# Newly repopulated spinal cord microglia exhibit a unique transcriptome and correlate with pain resolution

**DOI:** 10.1101/2022.12.20.521295

**Authors:** Lauren J. Donovan, Caldwell M. Bridges, Amy R. Nippert, Meng Wang, Shaogen Wu, Thomas E. Forman, Elena S. Haight, Nolan A. Huck, Sabrina F. Bond, Claire E. Jordan, Aysha S. Gardner, Ramesh V. Nair, Vivianne L. Tawfik

## Abstract

Microglia contribute to the initiation of pain, however, a translationally viable approach addressing how or when to modulate these cells remains elusive. We used a targeted, inducible, genetic microglial depletion strategy at both acute and acute-to-chronic transition phases in the clinically-relevant tibial fracture/casting pain model to determine the contribution of microglia to the initiation and maintenance of pain. We observed complete resolution of pain after transient microglial depletion at the acute-to-chronic phase, which coincided with the timeframe of full repopulation of microglia. These repopulated microglia were morphologically distinct from control microglia, signifying they may exhibit a unique transcriptome. RNA sequencing of repopulated spinal cord microglia identified genes of interest using weighted gene co-expression network analysis (WGCNA). We intersected these genes with a newly-generated single nuclei microglial dataset from human spinal cord dorsal horn and identified human-relevant genes that may ultimately promote pain resolution after injury. This work presents a novel approach to gene discovery in pain and provides comprehensive datasets for the development of future microglial-targeted therapeutics.

## MAIN TEXT

### Introduction

The transition to chronic pain is still poorly understood even though persistent pain affects one in three individuals.{Institute of Medicine, 2011 #73} Peripheral injury-induced neuronal hyperactivity leads to downstream engagement of microglia in the spinal cord,{Zhou, 2019 #1187} which contributes to the initiation of pain.{Haight, 2019 #1228;Ji, 2016 #1299} This has been confirmed in several preclinical pain models in which pharmacologic or genetic reversal of microglial activation {Gu, 2016 #1630;Tawfik, 2007 #608;Huck, 2021 #1614}or direct inhibition of spinal microglia with a Gi DREADD,{Yi, 2021 #1637} decreases nociceptive outcomes such as allodynia and thermal hyperalgesia. Additionally, depletion of microglia using either the immunotoxin Mac1-saporin,{Echeverry, 2017 #850;Sorge, 2015 #446} or transgenic Cx3CR1-Cre^ERT2-eYFP^;R26-iDTR^LSL^ mice{Peng, 2016 #863} prevents or reverses existing pain at limited time points after nerve injury. Importantly, after genetic depletion, microglia fully repopulate the CNS within 14 days.{Peng, 2016 #863;Parkhurst, 2013 #293} Therefore it is possible that either reactive microglia initiate the transition from acute-to-chronic pain, or that repopulated microglia actively resolve pain. The reversal of pain after transient depletion of microglia offers an opportunity to study the contribution of microglia in a pain-resolution context.

While microglial “signature genes’’ have been identified, transcriptomic studies highlight that microglia are heterogeneous cells that are phenotypically and genetically shaped by their microenvironment,{Lavin, 2014 #1642;Gosselin, 2014 #1643} particularly by their anatomic location.{De Biase, 2019 #1649} Importantly, the majority of these studies have been performed in distinct brain supraspinal regions{De Biase, 2017 #1650;Grabert, 2016 #759} with few to date undertaking transcriptomic analysis of microglia from the mouse spinal cord{Butovsky, 2014 #635;Fernandez-Zafra, 2019 #1077} and even fewer performing such analyses in human spinal cord tissue.{Tansley, 2022 #1751} Additionally, transcriptomic analyses of CNS tissue in disease states involving neuroinflammation, such as Alzheimer’s Disease,{Gosselin, 2017 #849} multiple sclerosis,{Masuda, 2019 #1640} and chronic pain,{Fernandez-Zafra, 2019 #1077} suggest heterogeneity in microglial responses to pathology. In-depth knowledge of the specific transcriptome of spinal cord microglia after pain-producing peripheral injury is therefore needed to identify markers for further targeted drug development for the treatment of chronic pain. Additionally, because these data are often collected in rodent models, the use of single-cell/nuclei sequencing data from human spinal cord is necessary to improve development of clinically translatable microglia-directed pain therapeutics.

In this study, we evaluated the temporal contribution of microglia to pain progression using targeted genetic microglia depletion with Cx3CR1-Cre^ERT2-eYFP^;R26-iDTR^LSL^ mice in the context of peripheral injury. We used the clinically-relevant tibial fracture/casting model of complex regional pain syndrome (CRPS) which has a distinctly timed acute-to-chronic pain transition. We discovered that microglial depletion/repopulation at the acute-to-chronic transition completely resolved pain and improved peripheral inflammation. We took advantage of this pain resolution context to establish transcriptomic signatures of repopulated spinal cord microglia at the point of full repopulation, which coincided with behavioral improvements in pain. Finally, we identified targets with human relevance by intersecting mouse genes of interest identified by weighted gene co-expression network analysis (WGCNA) with a newly generated human spinal cord single nuclei microglial transcriptome dataset.

## Results

### Morphologically distinct microglia repopulate the spinal cord 14 days after targeted genetic depletion

We utilized Cx3CR1-Cre^ERT2-eYFP^; R26-iDTR^LSL^ mice which allow for temporally controlled depletion of microglia, without depletion of other myeloid-lineage cells (ex. peripheral macrophages, Figure 1a). Tamoxifen was administered to express the diphtheria toxin receptor (DTR) in CX3CR1+ cells, then diphtheria toxin (DT) was administered 3-4 weeks later, giving time for peripheral macrophages to turn over, and no longer express the DTR. While these mice have been previously characterized,{Parkhurst, 2013 #293} we independently confirmed that the Cx3CR1-Cre^ERT2-eYFP^ mouse was tamoxifen dependent in a parallel study.{Huck, 2021 #1614} After DT-induced depletion, microglia repopulated to baseline numbers over the course of ∼14 days (Figure 1b-d), consistent with previous studies.{Bruttger, 2015 #430;Parkhurst, 2013 #293;Peng, 2016 #863} Interestingly, these repopulated microglia were morphologically distinct when compared to baseline control microglia, with fewer, shorter branches and an overall diminished complexity even at 14 days post-depletion/repopulation (Figure 1e-f).

**Figure 1.**
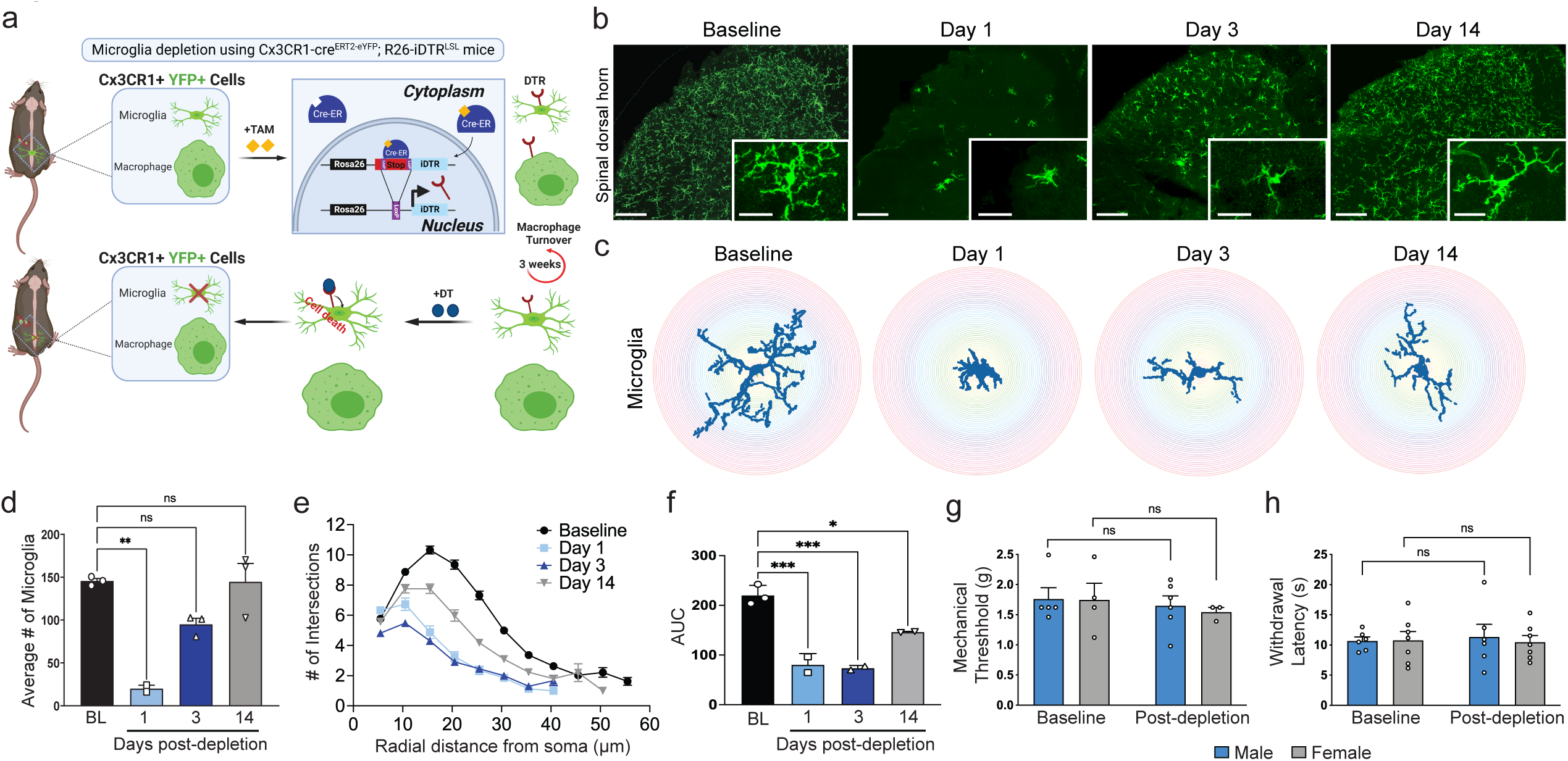
Cx3CR1-cre^ERT2^-eYFP; R26-iDTR^LSL^ mice exhibit significant loss of microglia after DT-induced depletion but these cells repopulate over time. a) Schematic representation of the strategy used to selectively deplete microglia, and no other myeloid-lineage cells, using the Cx3CR1-cre^ERT2-eYFP^;R26-iDTR^LSL^ mouse. Mice were injected with tamoxifen (TAM, 100 mg/kg daily x 5 days), allowing excision of the floxed STOP codon resulting in all Cx3CR1^+^ cells expressing the DTR. After 21-24 days, high-turnover macrophages no longer express DTR while slow-turnover microglia continue to express DTR. Treatment with diphtheria toxin (DT, 1000 ng daily x 3 days) resulted in selective death of microglia. b) Lumbar spinal cord sections from Cx3CR1-cre^ERT2-eYFP^; R26-iDTR^LSL^ mice taken at multiple time points before (baseline) or after DT injection demonstrate a significant loss of Iba1^+^ microglia in the dorsal horn at Day 1 post-DT by IHC. Scale bars 100 µm and 20 µm (inset). c) Representative Sholl analysis profiles of microglia demonstrate change in morphology over time following repopulation. d) Quantification of microglia number over time after depletion with DT. Four to five lumbar spinal cord dorsal horn sections were counted and averaged (n= 2-3 mice per group). **p<0.01, ****p<0.0001 by one-way ANOVA with Tukey’s post-hoc test. e) Analysis of intersections at a given radial distance from the soma demonstrates a drop in the number of intersections in repopulating microglia which recovers, though not quite to baseline, over 14 days. f) Area under the curve (AUC) for number of intersections by radial distance from soma demonstrates a time-dependent increase after microglial depletion. *p<0.05, ***p<0.001 by one-way ANOVA with Dunnett’s post-hoc test. Seventy-four to 185 microglia analyzed per time point with n = 2-3 mice per group. g) Microglial-depletion alone does not affect mechanical sensitivity or, h) latency to withdrawal on a 52.5°C hot plate in male or female Cx3CR1-cre^ERT2-eYFP^;R26-iDTR^LSL^ mice (n = 3-6/sex, comparison by unpaired t-test, in all cases p>0.05). ns, not significant. Data are represented as mean ± SEM.

### Microglial depletion at the acute-to-chronic transition drastically improves pain trajectory

We have previously demonstrated that the tibial fracture/casting model of CRPS results in signs of peripheral inflammation (paw edema, increased paw temperature) lasting up to 4 weeks post-injury. In contrast, mechanical paw sensitivity (allodynia) persists for at least 20 weeks.{Cropper, 2019 #1136;Huck, 2021 #1614} These findings suggest a peripheral-to-central (or acute-to-chronic) transition between 3-and 5-weeks post-injury. Importantly, microglia depletion alone did not affect baseline weight bearing, paw edema or paw temperature (Supplemental Figure 1a-c). Additionally, microglial depletion alone had no effect on measures of nociception, including unchanged mechanical sensitivity (Figure 1g) and hot plate latency (Figure 1h).

To establish the contribution of microglia to persistent pain, we first performed microglial depletion at the time of injury (day 0, acute phase) in the tibial fracture/casting model of CRPS (Supplemental Figure 1d). When microglia were depleted in this early/preventative paradigm, both male and female injured mice with microglial depletion (Injured-Depleted) demonstrated a significant but partial attenuation of allodynia over the course of the 9-week testing period, compared with sex-matched injured-only controls (Supplemental Figure 1e). The latency to withdrawal on a hot plate was significantly increased in females (*p* = 0.0304) but not males (*p* = 0.0780), in the Injured-Depleted groups (Supplemental Figure 1f-g). Both male and female Injured-Depleted mice improved their weight bearing on the injured hind paw at 4 weeks after injury (Supplemental Figure 1h-i), exhibited less edema (Supplemental Figure 1j-k), and less warmth (Supplemental Figure 1l-m) of the injured limb.

Due to the incomplete reversal of allodynia in the early/preventive phase depletion paradigm, as well as the clinical relevance of post-injury treatment, we next tested microglia depletion at 3 weeks post-fracture (acute-to-chronic phase, Figure 2a). Strikingly, we found that both male and female Injured-Depleted mice demonstrated a profound improvement in allodynia with a return to pre-injury baseline values over the course of the testing period (Figure 2b). The reversal of mechanical hypersensitivity was particularly remarkable as the mice remained allodynic immediately after DT treatment (when microglia were absent/depleted); then exhibited a progressively improving pain trajectory that temporally paralleled full spinal cord repopulation of microglia. We additionally observed an increase in hot plate latency in both male and female Injured-Depleted mice as well as improvement in weight bearing, edema, and warmth of the injured paw at 4 weeks in the Injured-Depleted mice (Figure 2c-d). These findings suggest that microglial repopulation, and not depletion itself, may trigger pain resolution and improvement in classic peripheral signs of CRPS (edema and temperature) in a post-injury/treatment paradigm.

**Figure 2.**
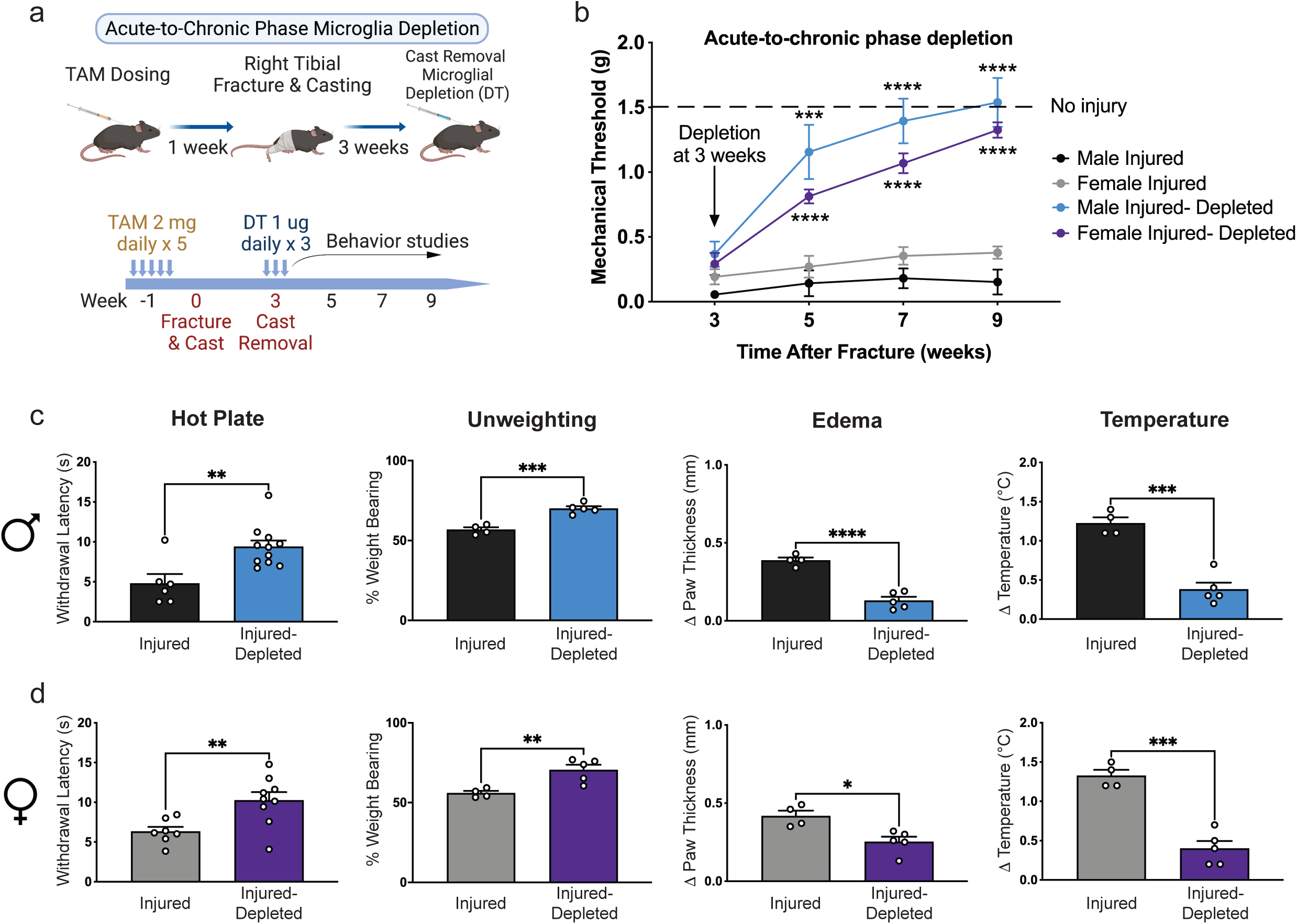
Microglial depletion at the acute-to-chronic phase results in a sustained change in pain trajectory after peripheral injury. a) Schematic of experimental timeline for acute-to-chronic phase depletion of microglia. b) Mechanical threshold is decreased at the time of cast removal (3 weeks) in all groups, however, microglial depletion results in sustained improvement in mechanical sensitivity in both males and females out to 9 weeks post-injury. n = 6-11 per group per sex. ***p<0.001, ****p<0.0001 by two-way ANOVA with Bonferroni’s post-hoc test. Microglial depletion at the acute-to-chronic phase additionally results in improvement in thermal sensitivity, weight bearing, edema and temperature changes in both males (c) and females (d) at 4 weeks post-injury. n = 6-11 per group per sex for hot plate, n = 4-5 per group per sex for unweighting, edema and temperature. *p<0.05, **p< 0.01, ***p<0.001 by two-tailed unpaired t-test. Data are represented as mean ± SEM.

### Repopulated microglia exhibit a distinct transcriptome that may contribute to pain resolution

In our model, morphologically distinct microglia repopulated by 14 days post-depletion, coincident with a sustained improvement in pain and peripheral inflammatory outcomes. To identify transcripts that may enable repopulated microglia to have a pro-resolution role in the context of peripheral injury, we FACS-isolated microglia from the lumbar spinal cord at 5 weeks post-injury (14 days post-microglia depletion/repopulation) for subsequent RNA sequencing from 4 groups: uninjured mice with resident microglia (Uninjured-Resident microglia), uninjured mice with repopulated microglia (Uninjured-Repopulated microglia), injured mice with resident microglia (Injured-Resident microglia) and injured mice with repopulated microglia (Injured-Repopulated microglia).

We sorted microglia as follows: live (SYTOX blue negative), CD19^-^CD3^-^ CD45^mid^CD11b^+^Cx3CR1-YFP^+^ (Supplemental Figure 2a). To first confirm the specificity of our isolation method for CNS microglia, we used the publicly available ImmGen “My GeneSet” program to cross-reference genes in our male RNAseq dataset with the ImmGen database of over 50 cell populations.{Heng, 2008 #770} The top 100 genes expressed by log2(cpm+1) in our datasets clearly defined a microglia-specific signature that could be differentiated from similar myeloid-lineage cells including blood monocytes (Supplemental Figure 2b). We next performed multidimensional scaling (MDS) and observed clustering of microglial gene profiles by treatment group, with microglia depletion/repopulation having the greatest effect (Supplemental Figure 2c-d), and these cells were distinct from Uninjured-Resident microglia.

Differential gene expression analysis using DESeq2 was performed on the microglial transcriptome for four comparisons by sex: Injured-Resident microglia vs. Uninjured-Resident microglia (effect of injury alone), Uninjured-Repopulated microglia vs. Uninjured-Resident microglia (effect of microglia depletion/repopulation alone), Injured-Repopulated microglia vs. Uninjured-Repopulated microglia (effect of injury in the context of microglia depletion/repopulation), Injured-Repopulated microglia vs. Injured-Resident microglia (effect of microglia depletion/repopulation in the context of injury). The analyses were performed at the transcript level for increased resolution and then transcripts were mapped to their corresponding genes for data representation, as indicated. For females we identified 34,865 transcripts which mapped to 13,377 unique genes. For males we identified 28,345 transcripts which mapped to 11,999 unique genes. Consistent with datasets from other pain models{Denk, 2016 #907}, the number of differentially expressed genes (DEGs) in sorted microglia in the injury alone context (Injured-Resident microglia vs Uninjured-Resident microglia) was limited. For females there were 4 upregulated DEGs and 8 downregulated DEGs, while for males there were 30 upregulated DEGs and 2 downregulated DEGs, none of which overlapped between sexes (Supplemental Table S1 and Extended Data 1). Also consistent with prior datasets,{Parkhurst, 2013 #293} microglial depletion/repopulation resulted in many DEGs (Uninjured-Repopulated microglia vs Uninjured-Resident microglia), indicating that repopulated microglia are distinctly different from microglia at baseline. Specifically, for females there were 251 upregulated DEGs and 375 downregulated DEGs, while for males there were 571 upregulated DEGs and 494 downregulated DEGs. In total, 99 upregulated DEGs and 72 downregulated DEGs overlapped between the sexes.

The effect of injury in the context of microglia depletion (Injured-Repopulated microglia vs Uninjured-Repopulated microglia) was sex-specific. For females there were 104 upregulated DEGs and 60 downregulated DEGs, while for males there were 92 upregulated DEGs and 7 downregulated DEGs. There were no upregulated DEGs that overlapped between the sexes, and just one shared downregulated DEG (*Slc4a7*) (Supplemental Table S1). The transcriptome of repopulated microglia therefore depends on the context (injury or no injury) in which they differentiate.

While the above comparisons provide a resource and confirm the validity of our dataset, to characterize microglia that repopulate in the context of peripheral injury but no longer contribute to sensitization (pain resolution context), we focused our subsequent analyses on the Injured-Repopulated microglia versus Injured-Resident microglia comparison. We identified numerous DEGs with several shared by both sexes. For females there were 293 upregulated DEGs and 338 downregulated DEGs, while for males there were 412 upregulated DEGs and 283 downregulated DEGs, (Figure 3a and Extended Data 1). In total, 112 upregulated DEGs and 72 downregulated DEGs overlapped between the sexes.

**Figure 3.**
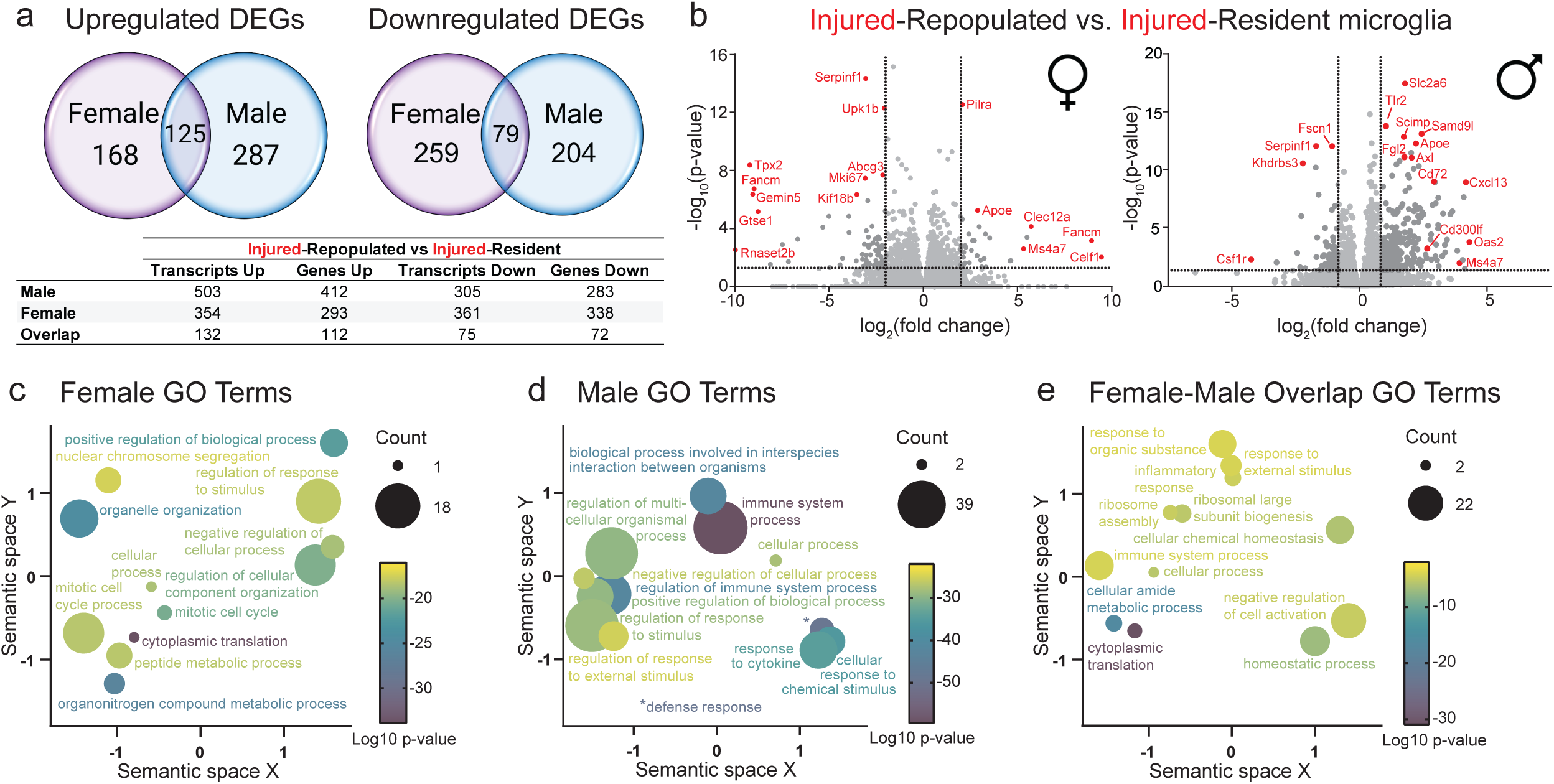
Differential gene expression analysis of female and male spinal cord microglia after peripheral injury with or without microglia depletion/repopulation identifies sex-specific and sex-independent signatures of pain recovery. a) Differentially expressed microglial transcripts (DETs) and genes (DEGs) were identified using DESeq2 analysis in both males and females. Overall males had more DEGs both up-and downregulated compared to females with 125 sex-independent/shared upregulated DEGs and 79 sex-independent/shared downregulated DEGs. b) Volcano plots depicting DEGs from injured-repopulated microglia compared to injured-resident microglia in females (left) and males (right). Highly regulated, sex-independent, and significant DEGs are labeled in red. Thresholds for the x-axis are set at |log_2_(FC)| >2.0 (female) and >0.83 (male). Threshold for the y-axis is set at -log_10_(p-value) > 1.3 for both sexes. Significant gene ontology (GO) terms for females (c) and males (d) based on adjusted p-value < 0.05 for injured-repopulated vs. injured-resident microglia DEGs. e) GO terms for the shared DEGs between male and female injured-repopulated vs. injured-resident microglia. Size of circle represents number of GO terms combined into the parent GO term shown. Color of circle represents log10 p-value.

To evaluate top regulated genes by sex in repopulated versus resident microglia with and without injury, we generated volcano plots to establish fold change to significance comparisons (Figure 3b and Supplemental Figure 3). We noted that *Apoe*, previously implicated in microglial phenotypic responses to injury,{Tansley, 2022 #1751} was highly upregulated in the Injured-Repopulated microglia in both sexes when compared to Injured-Resident microglia. We highlighted additional genes in volcano plots that were highly differentially expressed in Injured-Repopulated microglia or have been previously implicated in the context pain (Figure 3b, red text). Highly regulated female-specific genes included *Gtse1*, *Tpx2*, and *Upk1b (downregulated)*, as well as *Pilra*, *Clec12a*, and *Celf1* (upregulated). Highly regulated male-specific genes included *Csf1r*, *Khdrbs3*, *Fscn1* (all downregulated) as well as *Cd72, Cd300lf, Tlr2*, *Slc2a6*, *Axl* and *Cxcl13* (all upregulated). Highly regulated genes shared between the sexes included *Serpinf1* (downregulated), as well as *Apoe* and *Ms4a7* (upregulated) (Figure 3b, red text).

We next performed gene ontology (GO) term analysis on the DEGs from each comparison to identify biological pathways most strongly represented. In the female Injured-Repopulated microglia versus Injured-Resident microglia comparison, several GO terms related to cell cycle processes and regulation of cell component organization were represented, as well as “regulation of response to stimulus” (Figure 3c and Extended Data 2). In the male Injured-Repopulated microglia versus Injured-Resident microglia comparison, immune system processes were highly represented as were defense responses and responses to external stimuli/cytokines (Figure 3d and Extended Data 3). We next took all overlapping DEGs from male and female Injured-Repopulated microglia versus Injured-Resident microglia comparisons and ran GO analysis on these shared genes. (Figure 3e and Extended Data 4). While broad GO terms remained highly represented (cytoplasmic translation, homeostatic process, cellular process), we also identified GO terms related to the immune system including “inflammatory response”, “immune system processes”, and “response to external stimulus”, consistent with known microglial injury responses.{Huck, 2021 #1614} Taken together, this differential gene expression analysis highlights that repopulated microglia are transcriptionally responsive to injury with both sex-specific and sex-distinct gene changes while the behavioral phenotype (i.e. pain resolution) is similar between sexes.

### Construction of an unbiased gene co-expression network allows for discovery of novel genes contributing to pain resolution

While the DESeq2 analysis allowed us to identify individual differentially expressed genes, we next sought to discover biological processes and signaling pathways that better capture the functional difference between Injured-Resident microglia and the Injured-Repopulated microglia that contribute to pain resolution. To do this, we performed weighted gene co-expression network analysis (WGCNA), which clusters highly co-expressed genes into modules, without the use of *a priori* knowledge (Supplemental Figure 4). Importantly, gene expression in all samples are used in WGCNA, which allows for unbiased prediction of integral pathways associated with the cell states analyzed. Fifty-three unique modules were defined in females (Figure 4a, c) and 54 unique modules were defined in males (Figure 4b, c). Once these gene modules were constructed, the module eigengenes (ME), the first principal component of each module’s expression matrix, were then correlated by sex to each of the 4 trait comparisons to define significant module-trait relationships (Figure 4a-b, also see Extended Data 5 for a full list of correlation coefficients and *p*-values by trait).

**Figure 4.**
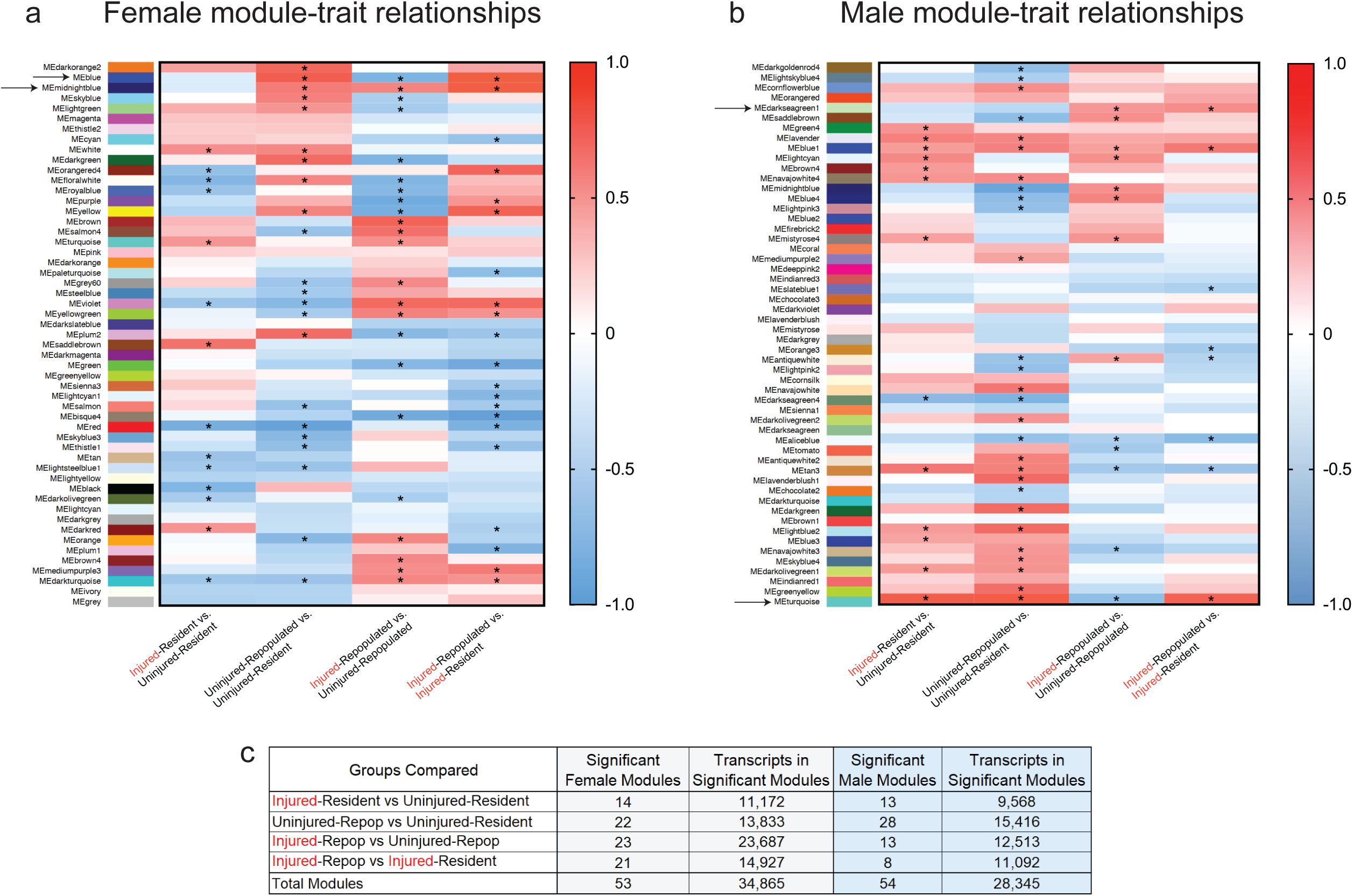
Weighted gene co-expression network analysis (WGCNA) reveals module-trait relationships that are distinct by sex and condition. WGCNA identified 53 unique modules in females (a) and 54 unique modules in males (b) which were then correlated with each of the 4 microglial conditions (“traits”). Module-trait relationships that had significant correlations are marked with an asterisk with heatmaps depicting the strength of the relationship (blue = negative correlation with the trait, red = positive correlation with the trait). c) The number of significant modules per sex as well as total number of transcripts within each module.

WGCNA identified networks of genes that characterized the pain resolution context with 21 significant modules in females and 8 significant modules in males in the Injured-Repopulated microglia to Injured-Resident microglia comparison (Figure 4c). To identify gene clusters strongly associated with pain resolution, we selected modules of interest for further analysis based on a combination of their absolute trait correlations across the 4 trait-comparisons (Figure 4a-b) and the number of differentially expressed transcripts (DETs) from the DESeq2 analysis represented in each module. We thus selected four modules [female: *blue* and *midnightblue;* male: *darkseagreen1* and *turquoise*], all of which had a strong correlation to the Injured-Repopulated versus Injured-Resident trait comparison (Figure 4a-b, arrows) and had high representation of DETs (Extended Data 6). To define highly represented biological processes and pathways, GO analysis and Ingenuity Pathway Analysis (IPA) were performed on all transcripts within each module, and genes highly impacting these pathways were further identified. Finally, we established the top 20 genes within each module with the highest gene significance vs. module membership, a measure of the strength of the correlation of the individual gene with the module eigengene (Figure 5). These genes are central to module construction because they correlate with a large number of genes within the module, acting as a major node (colored data points), and thus serve as natural candidates for further investigation (Figure 5).

**Figure 5.**
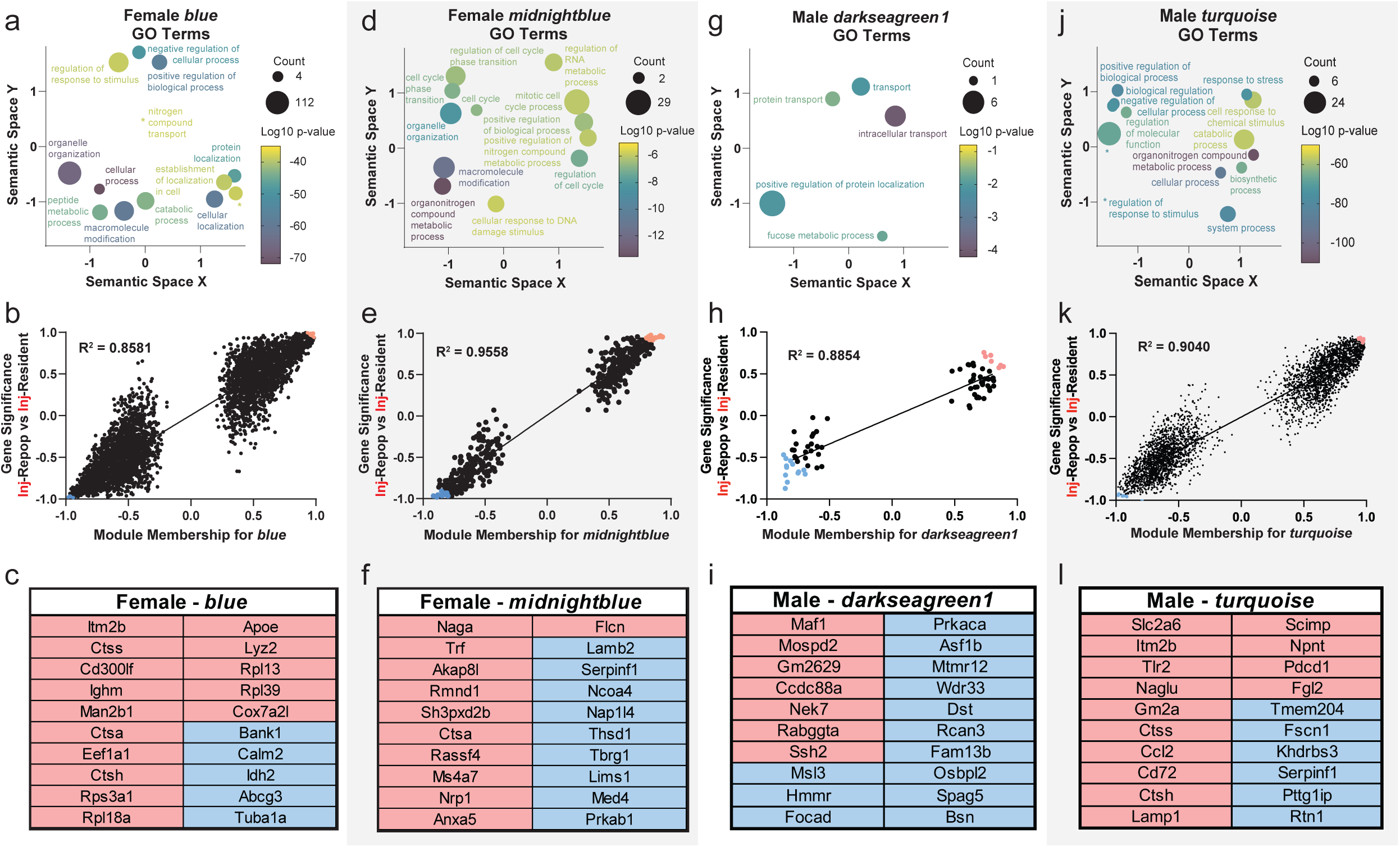
Modules of interest for each sex highlight networks of genes that may contribute to pain resolution. a) Gene Ontology (GO) terms identified for female module *blue*. b) Gene significance vs. module membership plot demonstrates correlation of member genes to the module eigengene. The 20 most extreme genes are highlighted either red for positive fold-change or blue for negative fold-change in the Injured-Repopulated microglia group versus Injured-Resident microglia group. c) List of top 20 most significant genes in gene significance versus module membership correlation for female module *blue*. d) GO terms for female module *midnightblue*. e) Gene significance vs. module membership plot for female module *midnightblue*. f) Top 20 most extreme genes for gene significance vs. module membership correlation for female module *midnightblue*. g) GO terms for male module *darkseagreen1*. h) Gene significance vs. module membership for male module *darkseagreen1*. i) Top 20 most extreme genes for gene significance vs. module membership correlation for male module *darkseagreen1*. j) GO terms for male module *turquoise*. k) Gene significance vs. module membership for male module *turquoise*. l) Top 20 most extreme genes for gene significance vs. module membership correlation for male module *turquoise*. Size of circle represents number of GO terms combined into the parent GO term shown. Color of circle represents log10 p-value.

For the female module containing the highest number of DETs, *blue,* we identified GO terms related to ‘regulation of response to stimulus’ consistent with identified IPA pathways ‘Clathrin-mediated Endocytosis Signaling’ (*Dnm2, Clta, Ap2b1*) which is upstream of actin polymerization. Additionally, the module is enriched for key genes in the ‘IL-12 signaling’ pathway (*Map2k3*, *Mapk12*, *Nfkb1*) involved in regulation of pro-inflammatory TNF-signaling as well as nitric oxide production.{Simpson, 2020 #1755} Interestingly, these key genes are all downregulated in our dataset, indicating reduced inflammation in Injured-Repopulated microglia. Regulation of plasma lipoprotein levels was also highly represented by GO analysis, and IPA included ‘LXR/RXR Activation’ (*Rxry*, *Apoe, Lyz2*), a pathway that affects lipoprotein clearance and cholesterol homeostasis (Figure 5a and Extended Data 7) {Holtman, 2017 #1819;Pena-Altamira, 2018 #1802;Jung, 2022 #1801}. For this module, 6 transcripts for the *Apoe* gene demonstrated positive significance (Figure 5b, c). Additionally, ‘Phagosome Maturation’ (*Ctss, Ctsh, Tuba1a*) was identified by IPA, an essential process following phagocytosis of debris/apoptotic cells (Extended Data 7). In line with this, a natural candidate identified in the *blue* module, *Cd300lf*, enhances phagocytosis {Borrego, 2013 #1781} (Figure 5b-c), and may enable Injured-Repopulated microglia to clear neuronal debris following injury.

In the female module *midnightblue*, GO terms identified represented broad categories such as ‘macromolecule modification’ and ‘positive regulation of biological processes’ (Figure 5d). IPA analysis highlighted more specific pathways and included ‘Acute Phase Response Signaling’ (*IL6st*, *Serpinf1*, *Trf*) with key genes downregulated and involved in inflammatory IL6-and TNF-signaling (Extended Data 7). In line with this, *Serpinf1*, an identified natural candidate downstream of IL6 signaling, is significantly downregulated (Figure 5e-f). Overall, this module represents reduced inflammatory pathway signaling which may contribute to pain resolution effects of Injured-Repopulated microglia.

In the male module *darkseagreen1*, significantly enriched GO terms (Figure 5g) were related to ‘regulation of protein transport’ and ‘protein localization’ with related enriched IPA pathways of ‘Calcium Signaling’ (*Arcan3*, *Prkaca*, *Asph*), which can change protein location within the cell {Clapham, 2007 #1806} and ‘Biosynthesis of Phosphatidylcholine and Choline’ (*Pcyt1a*), whose levels determine physical properties that affect fluidity and protein localization in the cell membrane (Figure 5g, Extended Data 7). One top 20 gene identified in *darkseagreen1* was *Prkaca*, which encodes the catalytic subunit of protein kinase A. Other GO terms enriched are related to pre-mRNA processing (*Wdr33*), which directly controls translation in the cell. Another top 20 gene found in this module was *Maf1*, a transcriptional regulator (Figure 5h, i). Overall, the module *darkseagreen1* emphasizes that these Injured-Repopulated microglia are tightly regulating protein expression, localization, and function in the cell, in particular the membrane.

In the male *turquoise* module, GO terms identified included ‘immune system process’ with IPA pathways such as ‘Neuroinflammation Signaling’ (*Irf7, Tlr2, Tnf, Cybb, Ccl2, Ccl5, IL12*) and ‘TREM1 Signaling’ (*Cd40, Itgax, Casp1, IL18, IL1*β). In addition, identical to the female *blue* module, male *turquoise* was enriched for the GO term ‘regulation of response to stimulus’ with related IPA pathways ‘LXR/RXR Activation’ (*Rxry*, *Apoe*, *Cd36, Il1rn*) and ‘Phagosome Maturation’ (*Cybb, Ctss, Ctsh, Lamp1, Rab7*) (Figure 5j and Extended Data 7). Specifically, the genes represented in this pathway are related to late phagosome maturation when lysosomal fusion forms the phagolysosome. One of the top 20 genes in this module was *Gm2a*, a gene encoding a co-factor of lysosomal enzymes. Further, two cathepsins (*Ctss* and *Ctsh*) with relevance to pain {Montague-Cardoso, 2020 #1810} were in the top 20 gene list in *turquoise (*Figure 5k-l). Ctss and Ctsh proteins are enriched in actively phagocytic cells, have lysosomal activity, and modify neuronal spine density extracellularly.{Hayashi, 2013 #1808} The *turquoise* module in males captures several biological functions similar to those identified in the female modules of interest, pointing towards a sex-independent phenotype of repopulating microglia active in phagocytosis and stress responses.

To compare the shift in gene expression in males and females captured by the WGCNA modules of interest, we plotted the fold change of transcripts from our selected modules in the Injured-Repopulated versus Injured-Resident microglia comparison (Supplemental Figure 5a) and observed a strong correlation between sexes (R^2^ = 0.6802). In contrast, when the fold change comparison was expanded to include all transcripts expressed in both sexes (Supplemental Figure 5b), there was no correlation (R^2^ = 0.0714). This suggests that WGCNA identified a shared transcriptomic shift in repopulated microglia in both males and females that may contribute to the pain resolution phenotype observed.

### Human dorsal horn spinal microglia exhibit heterogeneous cell states

Since our overall goal was to identify microglial targets for pain relief in humans, we next isolated human spinal cord dorsal horn microglia to cross-reference their transcriptomes with the generated mouse datasets. We performed single nuclei RNA sequencing (snRNAseq) on nuclei isolated from dorsal lumbar spinal cord from either a male or female post-mortem human donor with no known pain history (Supplemental Table 2). We isolated dorsal horn nuclei and enriched for microglia using negative selection sorting to provide an unbiased sample of microglia nuclei for sequencing (Figure 6a). The male and female snRNAseq datasets were then integrated and UMAP projection of the 6,591 nuclei revealed 7 clusters representing unique cell types. These clusters were defined using a human brain DropNc-seq dataset,{Habib, 2017 #1752} and included microglia, oligodendrocytes, oligodendrocyte precursor cells (OPCs), astrocytes, endothelial cells, neurons, and neuronal stem cells (NSCs) (Figure 6b and 6c). The negative selection strategy enriched microglial nuclei to ∼40% of the sample, compared to ∼3.7% collected from whole mouse spinal tissue in other studies{Matson, 2022 #1753} (Figure 6b). Microglial nuclei clusters expressed known marker genes described in previous studies including *ITGAM*, *CTSS*, *CSF1R* and *C1QA* (Figure 6d).{Tansley, 2022 #1751} One small nuclei cluster in the upper-left quadrant of the UMAP was determined to have high expression of T cell markers (and lacked microglial gene expression) and was thus eliminated before further sub-clustering of microglia (Figure 6c and d). A total of 2,316 microglial nuclei (dashed box in Figure 6c) were re-clustered using Seurat,{Stuart, 2019 #1756} revealing 6 distinct microglia sub-clusters based on unique gene expression profiles (Figure 6e, Supplemental Table 3, and Extended Data 8). GO analysis was then used to identify microglia sub-cluster cell states (Figure 6f and Extended Data 9). As expected from human post-mortem tissue, we detected a small cluster, cluster 6 (97 nuclei) of dying microglia with a top GO term “cell death”. We also detected a small cluster, cluster 5 (81 nuclei), that represented proliferative microglia with top terms such as “cell cycle”, “DNA metabolic processes”, and “chromosome organization”. Notably, cluster 5 expressed cluster-unique gene *MKI67* (p-value adj = 6.02E-134), the gene encoding Ki-67, as well as *NDC80*, *TOP2A* and *POLQ*, each prominent markers of proliferation (Supplemental Figure 6a and Extended Data 8). We identified an additional sub-cluster, cluster 4 (81 nuclei), with a signature similar to reactive microglia with GO terms such as “cell activation”, “regulation of cell motility”, and a cluster-unique term “inflammatory process”, with key genes significantly expressed such as *TLR2* and *CYBB* (also known as *NOX2*), both potent neuroinflammatory-associated genes expressed by microglia{Yang, 2020 #1754;Simpson, 2020 #1755} (Supplemental Figure 6b). Additionally, a top differentially expressed gene in cluster 4 was AC083837.1 (p-value adj = 9.96E-105), a novel long noncoding RNA induced in the human microglial cell line (HMC3) in response to LPS treatment {Baek, 2022 #1820} (Supplemental Figure 6b). Importantly, we further identified a large nuclei cluster that represented homeostatic microglia (cluster 1, 730 nuclei), with cluster-unique terms “myeloid cell homeostasis” and “regulation of supramolecular fiber organization” as well as “regulation of cell adhesion” and “cell junction organization”. Anti-inflammatory or homeostatic genes such as *TGFBI*, *ITGAX*, and *IGF1R*, were significantly expressed with positive fold-changes, while inflammatory-associated genes, such as *TLR2* and *CYBB,* had negative fold-changes in homeostatic cluster 1 compared to other clusters (Supplemental Figure 6b-c and Extended Data 8). The final two clusters (cluster 2, 857 nuclei; cluster 3, 470 nuclei) had both homeostatic and reactive features with terms shared by homeostatic cluster 1, such as “immune system process” and “regulation of cell adhesion”, in addition to terms shared by clusters 4 and 6, such as “cell activation” and “cell death” (Figure 6f).

**Figure 6.**
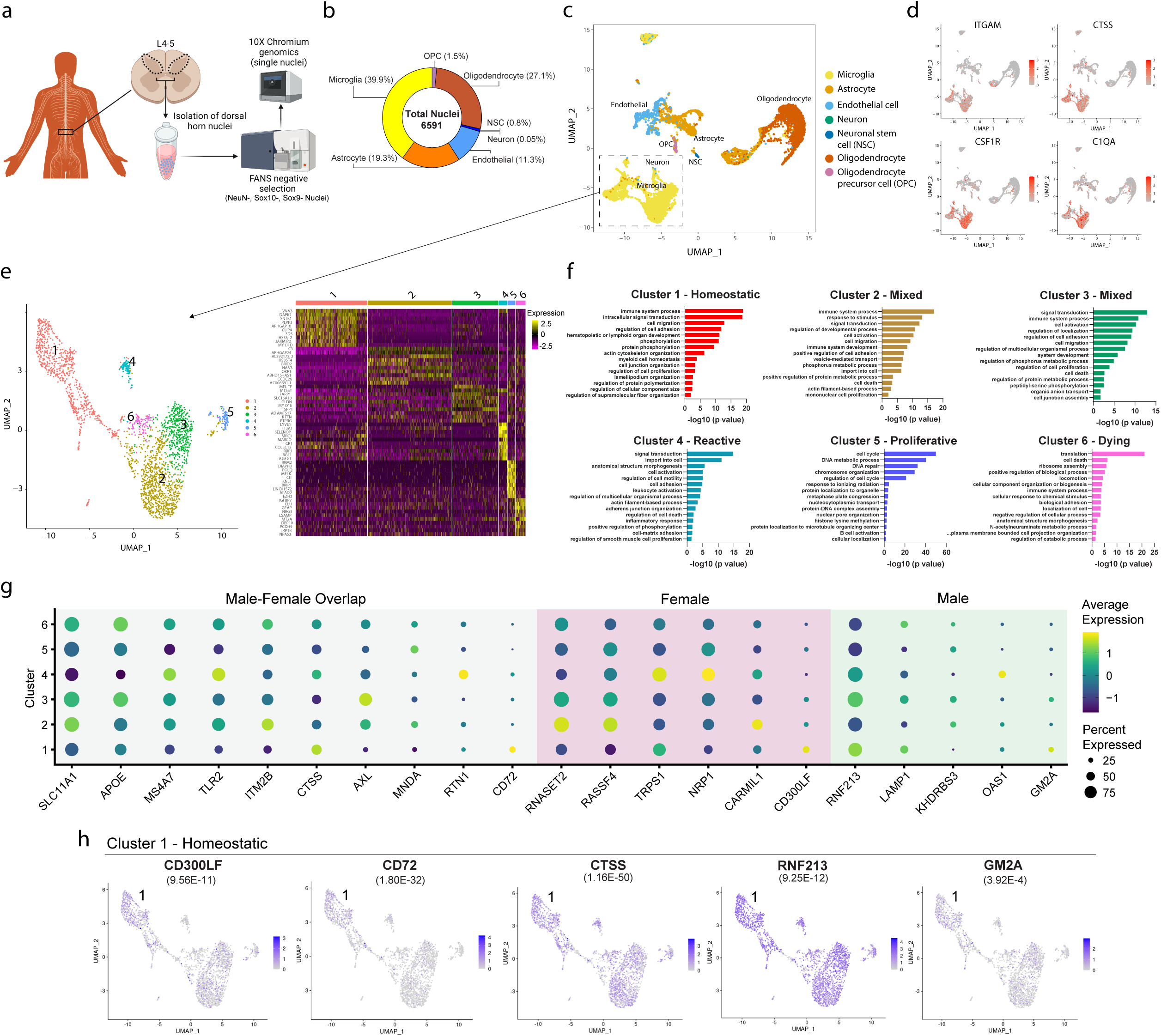
Single-nuclei RNA-sequencing (snRNA-seq) of human microglia uncovers gene targets identified in repopulated microglia. a) Diagram depicting experimental flow of isolation and enrichment of microglial nuclei from human spinal cord dorsal horn grey matter and subsequent 10X Chromium sequencing. b) Percentages of cell-type specific nuclei captured and sequenced, with microglia enriched to 40% of total nuclei. c) Uniform manifold approximation and projection (UMAP) projection showing distribution of human spinal cord single nuclei clusters by cell-type. Dashed box indicates final microglial clusters that were further sub-clustered in panel *e*. d) Microglial-specific cell clusters are identified using known microglia gene markers. e) Six distinct microglial nuclei clusters were identified after sub-clustering using Seurat. f) Gene ontology (GO) analysis showing top 15 GO terms that capture differential microglia cell states. g) Dot plot of selected top mouse genes that were significantly differentially expressed between human microglial subclusters (cluster 1-6). Genes that overlap in male/female mouse datasets shown in the grey zone, female-specific genes shown in the purple zone, and male-specific genes in green zone. h) UMAP plots showing genes significantly highly expressed in the human homeostatic cluster 1, also upregulated in injured-repopulated mouse microglia. Adjusted p-value from differential cluster-expression analysis shown in parentheses below gene name. Scale bar represents relative gene expression.

### Identification of human-relevant homeostatic gene targets detected in repopulated mouse microglia

To identify human-relevant microglia-expressed genes found in our Injured-Repopulated microglia vs Injured-Resident microglia mouse dataset, we first selected mouse transcripts that met a DESeq2 cutoff of -log10(p-adjusted) > 1.3. We then ranked the transcripts in descending order of their differential expression fold-change and adjusted p-value and removed transcripts that weren’t within our top WGCNA modules of interest. We next categorized these top mouse transcripts as having either sex-independent or sex-specific expression. Lastly, we mapped the transcripts to their corresponding gene and the resulting mouse gene list contained 12 top female genes, 29 top male genes, and 30 overlap genes (Supplemental Table 4). These mouse genes were then screened across the human microglial nuclei clusters to identify cluster-specific expression patterns. Twenty-one of these mouse genes were significantly differentially expressed across the human microglia clusters (p-value adj <0.05) (Figure 6g). We identified one gene, *KHDRBS3* (also known as *SLM2*), that had significantly lower expression, specifically in homeostatic human cluster 1, and was also down-regulated in mouse Injured-Repopulated microglia. Additionally, five genes (*CTSS*, *CD72, CD300LF, RNF213*, and *GM2A,*) had significantly higher expression in homeostatic human cluster 1, and no other cluster (Figure 6g-h). Interestingly, all these genes were upregulated in mouse Injured-Repopulated microglia (Supplemental Table 4), indicating possible involvement in homeostatic functions in humans and thus may serve as relevant microglial targets to promote spinal cord homeostasis and pain resolution.

## Discussion

In this study, we established the importance of microglia at multiple time points after pain-producing peripheral injury, identified transcriptomic signatures of spinal cord microglia associated with a pain-resolved state, and screened top gene candidates with those expressed by microglial subclusters obtained from human spinal cord transcriptional profiling. We applied a genetic depletion strategy using Cx3CR1-Cre^ERT2-eYFP^;R26-iDTR^LSL^ mice to temporally and selectively deplete microglia without affecting peripheral circulating monocytes/macrophages.{Parkhurst, 2013 #293;Huck, 2021 #1614} We find that depletion of microglia at the acute-to-chronic pain transition (at 3-week post-injury) is sufficient to resolve existing allodynia and thermal hyperalgesia in both sexes. Importantly, the onset of pain resolution coincided with spinal cord microglial repopulation, not microglial depletion. To identify a potential “pro-resolution” gene signature in these repopulated spinal cord microglia, we performed RNA sequencing followed by bioinformatic analysis using DESeq2 and WGCNA. WGCNA is a powerful technique that has historically been applied to population-level studies of publicly available datasets.{Neidlin, 2019 #1800;Topouza, 2022 #1799} Our study is unique in applying WGCNA in the context of pain transcriptomic analysis of microglia. Using this in-depth, unbiased approach, we identify both sex-dependent and independent microglial transcriptional signatures related to the uninjured, injured, or pain-resolved states. Given the limitations of using mice as a model for human disease, we next generated a snRNA-seq dataset from human dorsal horn lumbar spinal cord microglial nuclei. We characterized heterogeneity of microglia in this anatomically limited region, including 6 subclusters with unique transcriptomes, some of which shared similarities with previously identified subpopulations in brain.{Chen, 2021 #1788} Finally, we cross-referenced candidate mouse microglial genes linked to the pain-resolved state with this human snRNA-seq dataset and identified genes of high translational interest based on their sub-cluster expression profiles.

Microglia are associated with the acute phase/initiation of pain after peripheral injury{Zhou, 2019 #1187}; our data using genetic microglial depletion suggests they may also contribute to the transition from acute-to-chronic pain, and perhaps even to pain resolution. Previous studies have demonstrated that microglial activation is necessary{Yi, 2021 #1637} and sufficient{Yi, 2021 #1636} to initiate acute pain. Microglial depletion studies using pharmacologic or genetic tools show that microglia also contribute to the initiation of pain progression in models of nerve injury{Echeverry, 2017 #850;Sorge, 2015 #446;Peng, 2016 #863}. We find that preventive depletion of microglia (at the time of injury) was insufficient to completely reverse allodynia. However, the extent of improvement was similar in magnitude to that observed after early microglial depletion in a high-frequency stimulation model of pain{Zhou, 2019 #1187} and in spinal nerve transection.{Peng, 2016 #863} This lack of full recovery may be due to nociceptor hyperactivity or high levels of acute nociceptor-derived inflammatory signals acting on the spinal cord environment immediately post-injury.{Nieto, 2015 #1787}

Of note, consistent with our own findings, several studies{Peng, 2016 #863;Zhou, 2019 #1187;Mousseau, 2018 #847;Kohno, 2022 #1774} reported equivalent improvements in pain outcomes in both sexes with microglial manipulation. Taken together, this suggests that microglia themselves are unlikely to be sex-specific contributors to pain progression, but rather microglial expressed receptors such as TLR4{Huck, 2021 #1614;Sorge, 2011 #469} or chemokines such as CCR2{Zhou, 2019 #1187} may be sex-specific.

The current work temporally extends the involvement of microglia to the acute-to-chronic phase, with microglial depletion at 3 weeks post-injury significantly and persistently improving pain trajectory to baseline levels. In agreement, myeloid cell depletion with the CSF1 inhibitor PLX5622 at 28-days post-peripheral nerve injury reversed mechanical and cold allodynia.{Lee, 2018 #1792} In contrast, Peng et al.{Peng, 2016 #863} demonstrated that early depletion of microglia delayed development of allodynia, while late depletion of microglia only transiently improved allodynia. The reason for this discrepancy may relate to the type of pain model used (direct nerve injury vs. fracture/immobilization), or exact timing of microglial depletion (week 1 vs 3 post-injury). In contrast to the commonly used nerve injury models,{Decosterd, 2000 #774;Kim, 1992 #1797} the tibial/fracture model of CRPS exhibits a clearly timed transition from acute-to-chronic pain, and triggers both inflammatory and neuropathic mechanisms, mimicking the clinically relevant human condition.{Cropper, 2019 #1136;Haight, 2020 #1433}

In our study, pain resolution after microglial depletion at the acute-to-chronic phase coincided with the timeframe of complete repopulation of CNS microglia. This outcome suggests that repopulated microglia may re-establish homeostasis within the pain circuit. Indeed, newly repopulated microglia can function to repair synapses in the hippocampus as well as promote recovery following brain injury{Basilico, 2022 #1767;Rice, 2017 #1768}. These cells also retain the ability to respond to environmental cues and our WGCNA analysis highlights that they remain highly active cells engaged in surveillance and phagocytosis.{Yao, 2016 #1652;Huang, 2018 #1749} The possibility that repopulated microglia exhibit a phenotype that is distinct from resident microglia is further supported by transcriptome analyses from prior published work showing clear differences between repopulated and “control” brain microglia.{Bruttger, 2015 #430;Huang, 2018 #1749} One major finding of the current work is that the environment (either injured or uninjured) in which microglia repopulate also influences microglial phenotype. This is perhaps unsurprising given that microglia are dynamic, multidimensional cells constantly surveying their environment and responding to stimuli with a range of state changes.{Paolicelli, 2022 #1776;Hammond, 2019 #1785} An urgent question remains regarding whether these heterogenous microglial states may have functional significance in disease progression or resolution, in particular for pain.

Since microglia repopulation did not result in re-sensitization of the periphery, and repopulated microglia were morphologically distinct from control microglia, our model allowed for identification of genes with potential to contribute to pain resolution. The intriguing possibility that specific populations of microglia may contribute to pain resolution is supported by a seminal study characterizing CD11c+ spinal cord microglia as necessary and sufficient to the resolution of pain after peripheral nerve injury in both sexes.{Kohno, 2022 #1774} Aside from high expression of *Itgax*, *Igf1* and *Apoe*, this antinociceptive subpopulation also highly expressed the receptor tyrosine kinase, *Axl*. Importantly, we also find upregulation of *Apoe* and *Axl* in both male and female repopulated microglia isolated from the pain resolution group, however, in a model of neuropathic pain *Apoe* was elevated in resident microglia after injury{Tansley, 2022 #1751}. Furthermore, *Apoe* is upregulated in disease-associated microglia in other contexts {Keren-Shaul, 2017 #957;Absinta, 2021 #1789}. The elevated level of *Apoe* in a diversity of microglial states, spanning reactive to resolving, suggests the gene may in fact be a non-specific marker for highly active rather than disease-promoting microglia, especially in the context of pain. In support of this interpretation, we detected widespread expression of *APOE* in our human microglial nuclei clusters, a finding supported by a recent human spinal cord sequencing study.{Yadav, 2022 #1798} Additional studies directly testing the function of *Apoe* in microglia in the context of pain will be necessary to implicate it as a pain-regulating gene.

For translatable therapies to be developed, it is necessary to validate mouse-identified gene targets using human tissue samples. To address this, we profiled single nuclei of human spinal microglia taken specifically from the lumbar grey matter (dorsal horn). We thus focused on microglia in a pain-relevant region, which is crucial given that gene signatures and cellular states are defined by microglial environment.{Gosselin, 2017 #849;Yvanka de Soysa, 2022 #1777;Xuan, 2019 #1778} We performed negative selection (removing astrocytes, neurons, and oligodendrocytes) to enrich the sample 10-fold for microglia, while also capturing those not expressing canonical positive-selection markers (ex. Iba1 or Cx3CR1). This enhanced our ability to characterize human microglial states, which are known to be more heterogeneous compared to other species including the mouse.{Geirsdottir, 2019 #1779} Using this region-specific, negative-selection approach, we uncovered 6 microglial clusters that were spatially distinct by UMAP and displayed unique gene expression patterns. Of note, we observed one large spatially distinct cluster by UMAP (cluster 1) that contained homeostatic gene expression signatures and lacked reactive ones.

To concentrate our gene identification to those with human-relevance, we screened top candidate mouse genes for expression within human microglia clusters. We identified several genes significantly upregulated in repopulated mouse microglia from the pain resolution group, also significantly, and specifically, expressed by human homeostatic cluster 1. *CD300LF* (the human conserved ortholog of the mouse gene *cd300f* {Clark, 2009 #1782}) encodes a member of the CD300 cell surface glycoprotein family that functions to positively regulate phagocytosis of apoptotic cells, this function is increased in the spinal dorsal horn microglia subpopulation shown to promote pain resolution following injury.{Borrego, 2013 #1781;Kohno, 2022 #1774} Specifically, *CD300LF* is an inhibitory receptor for myeloid cells that negatively regulates TLR signaling via MYD88.{Kim, 2012 #1780} In addition, over-expression of *CD300LF* has a neuroprotective effect after acute brain injury,{Peluffo, 2012 #1783} and microglia isolated from mice resistant to cerebral malaria express high levels of *cd300lf* and inhibit inflammation.{Keswani, 2020 #1784} Another gene identified was *CD72*, whose protein product is a receptor for semaphorin 4D and contributes to microglial-mediated CNS inflammation.{Kumanogoh, 2000 #1794;Okuno, 2010 #1793} Lastly, we identified *GM2A*, which encodes a small, secretable glycolipid transport protein and stimulates the degradation of ganglioside GM2 in the lysosome. Elevated levels of GM2A in culture reduce neurite extension and neuronal firing rate, which may be beneficial in the context of pain and dorsal horn circuitry.{Hsieh, 2022 #1795}

In this study, we took advantage of a pain resolution context to profile repopulated spinal cord microglia and identify genes that may promote homeostatic features of microglia, and ultimately promote pain recovery after injury. Future studies will evaluate whether these repopulated microglia directly contribute to the resolution of pain, similar to previously described CD11c+ microglia,{Kohno, 2022 #1774} or whether simply the absence of reactive microglia in a short time window prevents chronic pain in our depletion studies. Our findings, that repopulated microglia are transcriptomically distinct from resident microglia after injury, suggest they contribute in a unique manner to the spinal cord circuit environment. Interrogating the individual genes identified in this study which overlap with human microglia subpopulations, many of which have no known function in microglia, will be an exciting next step to confirm their potential as therapeutic targets for analgesic development.

## Acknowledgments

Flow cytometry and Fluorescent Activated Cell Sorting (FACS) was done with instruments in the Palo Alto VA Flow Cytometry Core, which is supported by the US Department of Veterans Affairs (VA), Palo Alto Veterans Institute for Research (PAVIR), and the National Institutes of Health (NIH). Additional cell sorting/flow cytometry analysis for this project was done on instruments in the Stanford Shared FACS Facility. The authors wish to thank Dr. Lianna Bonnano, Dr. Corey Cain, Dr. Alex Lee and Pratima Nallagatla for assistance with FACS and initial bioinformatic analyses. We thank Janelle Siliezar-Doyle and Dylan Mayanja for contributions to initial behavioral analyses. We are also indebted to Dr. Long-Jun Wu and Dr. Min-Hee Yi for training on microglial Sholl analysis. We thank Dr. Brendan Hasz for building our publicly-available website for data sharing and Dr. Arkady Khoutorsky for comments on early versions of this work. This work utilized computing resources and computational and bioinformatics services provided by the Stanford Genetics Bioinformatics Service Center (GBSC).

## Funding

National Institutes of Health (NIH) grant R35GM137906 (VLT)

National Institutes of Health grant T32DA035165 (LJD)

Rita Allen Foundation Scholars Program Fund (VLT)

Philanthropic donation from the Duan Family (VLT)

## Author contributions

Conceptualization: LJD, VLT

Methodology: TEF, VLT

Formal analysis: LJD, CMB, ARN, MW, SW, CEJ, ASG, RVN, VLT

Investigation: LJD, ARN, SW, TEF, ESH, NAH, VLT

Writing—original draft: LJD, CMB, ARN, VLT

Writing—review & editing: SW, ESH, NAH, SFB, CEJ, ASG, RVN, VLT

Visualization: LJD, CMB, ARN, VLT

Supervision: RVN, VLT

Project administration: VLT Funding acquisition: LJD, VLT

## Data and materials availability

Transcriptomic data are included as Extended Data Tables. All transcriptomic data have been deposited as raw FASTQ files in GEO database accession number (in process). We have created two publicly-available, browsable interfaces as a resource for the field https://tawfik-lab-repopulated-microglia-transcrip-streamlit-app-xsbwen.streamlit.app, which includes all transcriptomic bulk-seq mouse spinal cord microglia data and https://tsomics.shinyapps.io/visualization_MGcells/ which includes the human microglial nuclei sequencing data. All other data supporting the findings of this study are available from the corresponding author upon reasonable request.

## Materials and Methods

### Animals

Adult male and female mice 10-12 weeks old at the start of the experiments were housed 2-5 per cage maintained on a 12-hour light/dark cycle in a temperature-controlled environment with *ad libitum* access to food and water. Male mice weighed approximately 25 g at the start of the study, and female mice weighed approximately 20 g at the start of the study. Mice used in this study: wild type C57BL/6J mice (Jax stock #00664), CX_3_CR1^CreERT2-EYFP^ (Jax stock #021160),{Parkhurst, 2013 #293} Rosa26-loxP-stop-DTR (R26^iDTR^, Jax stock #007900). To specifically and conditionally ablate central CX_3_CR1 cells (microglial depletion), we crossed homozygous CX_3_CR1^CreERT2-EYFP^ with homozygous R26^iDTR/iDTR^ mice to generate double heterozygous Cx3CR1-Cre^ERT2-eYFP^;R26-iDTR^LSL^ mice. Mouse genotypes from tail samples were determined using real time RT-PCR with specific probes designed for each gene (Transnetyx, Cordova, TN). Control subjects for all behavioral studies were mice without the R26^iDTR^ allele: Cx3CR1-Cre^ERT2-eYFP^; R26-iDTR^+/+^ that received tamoxifen (TAM) and diphtheria toxin (DT) with the same dosing and administration schedule as experimental mice.

### Drugs and Route of Administration

Tamoxifen (Sigma, #T5648) was dissolved in corn oil at a concentration of 25 mg/mL. Mice were injected intraperitoneally (i.p.) at 100 mg/kg daily for 5 days. Diphtheria toxin (DT) from *Corynebacterium diphtheriae* (Sigma, #D0564-1MG) was dissolved in sterile water to a concentration of 10 µg/mL and 0.1 mL (1000 ng) was administered i.p. daily for 3 days.

### Tibial fracture/casting model of complex regional pain syndrome (CRPS)

Mice were anesthetized with isoflurane and underwent a closed right distal tibia fracture followed by casting. The right hind limb was wrapped in gauze and a hemostat was used to make a closed fracture of the distal tibia. The hind limb was then wrapped in casting tape (ScotchCast™ Plus) from the metatarsals of the hind paw up to a spica formed around the abdomen to ensure that the cast did not slip off. The cast over the paw was applied only to the plantar surface with a window left open over the dorsum of the paw and ankle to prevent constriction when post-fracture edema developed. Mice were inspected throughout the post-operative period of cast immobilization to ensure that the cast was properly positioned. At 3 weeks post-fracture, mice were briefly anesthetized, and casts were removed with cast shears. For behavioral assessment, mice were tested beginning three days after cast removal at 3 weeks until 9 weeks post-fracture, as indicated in each section below. CRPS model generation and behavioral testing were conducted following well established methods for evaluating mouse behavior in the tibial fracture-casting model of CRPS.{Birklein, 2018 #1183;Guo, 2014 #332}

### Behavioral testing

To ensure rigor in our findings and avoid the contribution of experimenter sex to our behavioral data, experimenters were female or there was a female scientist’s lab coat in the room during acclimation and testing.{Sorge, 2014 #955} *In vivo* behavioral testing was performed in a blinded fashion. All testing was conducted between 7:00 am – 1:00 pm in an isolated, temperature-and light-controlled room. Mice were acclimated for 30 – 60 minutes in the testing environment within custom clear plastic cylinders (4” D) on a raised metal mesh platform (24” H). Mice were randomly placed in a cylinder; after testing, mouse identification numbers were recorded on the data sheet.

### Mechanical nociception assays

To evaluate mechanical reflexive hypersensitivity, we used a logarithmically increasing set of 8 von Frey filaments (Stoelting), ranging in gram force from 0.007 to 6.0 g. These were applied perpendicular to the plantar hind paw with sufficient force to cause a slight bending of the filament. A positive response was characterized as a rapid withdrawal of the paw away from the stimulus filament within 4 s. Using the up-down statistical method,{Chaplan, 1994 #966} the 50% withdrawal mechanical threshold scores were calculated for each mouse and then averaged across the experimental groups. Mechanical nociception testing was performed at Weeks 3, 5, 7, and 9 post-fracture.

### Paw Edema, Unweighting and Temperature Measurements

Hind paw edema was determined by measuring the hind paw dorsal-ventral thickness over the midpoint of the third metatarsal with a LIMAB laser measurement sensor (LIMAB, Goteborg, Sweden) while the mouse was briefly anesthetized with isoflurane. Temperature and hind paw thickness data were analyzed as the difference between the fracture side and the contralateral intact side and averaged across experimental groups. Paw edema was measured at 4 weeks post-fracture.

An incapacitance device (IITC Inc Life Science, Woodland Hills, CA) was used to measure hind paw unweighting. Mice were manually held in a vertical position over the apparatus with the hind paws resting on separate metal scale plates, and the entire weight of the mouse was supported on the hind paws. The duration of each measurement was 6 seconds, and 6 consecutive measurements were taken at 60-second intervals. Six readings were averaged to calculate the bilateral hind paw weight-bearing values. Unweighting was measured at baseline and then again at 4 weeks post-fracture. Data was analyzed as a ratio between the right hind paw weight divided by the sum of right and left hind paw values [2R/(R + L)] × 100%).{Li, 2014 #358}

The temperature of the hind paw was measured using a fine-gauge thermocouple wire (Omega, Stamford, CT). Temperature testing was performed over the hind paw dorsal skin between the first and second metatarsals (medial), the second and third metatarsals (central), and the fourth and fifth metatarsals (lateral). The measurements for each hind paw were averaged for the mean paw temperature. Data were expressed as the average difference between the ipsi-and contralateral hind paw within an experimental group. Paw temperature was measured at 4 weeks post-fracture.

### Thermal nociception assays

To evaluate thermal-induced reflexive responses, we used the hotplate test (plate temperature was set to 52.5°C). Mice were placed on the plate and the latency (seconds) to the first appearance of a reflex response (flinch) was recorded as a positive reflex withdrawal response. A maximal cut-off of 45 s was set to prevent tissue damage. Only one exposure to the hotplate was applied on a given testing session to prevent behavioral sensitization that can result from multiple noxious exposures. Measurements were made at 5 weeks post-fracture.

### Immunohistochemistry

Mice (12 – 30 weeks) were transcardially perfused with 10% formalin in PBS. The spinal cord (lumbar cord L3 – L5 segments) were dissected from the mice and cryoprotected in 30% sucrose in PBS and frozen in O.C.T. (Sakura Finetek, Inc.). Spinal cord sections (40 μm) were prepared using a cryostat (Leica Biosystems) and incubated in blocking solution (5% normal donkey serum and 0.3% Triton X-100 in PBS) for 1 h at room temperature followed by incubation with primary antibodies at 4°C, overnight. The following primary antibodies were used: rat anti-CD11b (Bio-Rad, #MCA711G, 1:500), and rabbit anti-Iba1 (Wako, #019-19741, 1:500). Tissues were washed with 1X PBS 3 times and sections were incubated with appropriate secondary antibody conjugated to AlexaFluor in 1% normal donkey serum, 0.3% Triton X-100, and 1X PBS for 2 h at room temperature. Sections were mounted with Fluoromount G with DAPI medium (ThermoFisher, #00-4959-52). Images were collected with a Keyence BZ-X800 fluorescent microscope (Keyence) using the sectioning module to remove non-focused light using 20x (NA 0.75) or 60x (NA 0.95) objectives. Four to five lumbar spinal cord dorsal horn sections were counted and averaged for 2-3 mice per group.

### Sholl analysis for microglial morphology

Lumbar spinal cord was sliced on a cryostat at 40 μm and microglia were labeled by immunohistochemistry using the rabbit anti-Iba1 antibody (Wako, #019-19 741, 1:500). Dorsal horn localized microglia were imaged using a Keyence BZ-X800 fluorescent microscope (Keyence) using the sectioning module (1D Slit method) to remove non-focused light using a 40X objective. Each z-stack was taken at the same exposure using 43 slices with a 0.3 μm step size. Images were imported into ImageJ/FIJI and maximum intensity projections of dorsal horn images were generated. Region of interest (ROI) was selected around individual microglia. Unsharp Mask was applied using set radius to 3 pixels and weight of 0.6 followed by the de-speckle tool was to remove non-specific background. The images were made binary and ‘remove outliers’ was used at threshold of 50. The Eraser tool was then used to manually eliminate Iba1+ processes within the frame that were clearly not associated with the microglial cell analyzed.

Between 37-92 microglia per animal per time-point (Baseline: 185 microglia, Day 1: 74 microglia, Day 3: 117 microglia, Day 14: 87 microglia) were analyzed by Sholl analysis plugin in ImageJ/FIJI with a start radius of 0.5 μm and a step size of 1 μm.

### Statistics for behavioral paradigms and Sholl analysis

Cohort sizes were determined based on historical data from our laboratory using a power analysis to provide >80% power to discover 25% differences with p<0.05 between groups to require a minimum of 4 animals per group for all behavioral outcomes, and 2 animals per group for Sholl analyses with >30 microglia counted per animal per time-point. All experiments were randomized by cage and performed by a blinded researcher. Researchers remained blinded throughout histological, biochemical, and behavioral assessments. Groups were unblinded at the end of each experiment before statistical analysis. Data are expressed as the mean ± SEM. Statistical analysis was performed using GraphPad Prism version 8.4.1 (GraphPad Software) or R, as described in Methods. Data were analyzed using a Student’s t tests, or ordinary one-way with Dunnett’s or Tukey’s post-hoc test, or two-way analysis of variance with a Bonferroni post-hoc test, as indicated in the main text or figure captions, as appropriate, with complete statistical analyses detailed in Table. The “n” for each individual experiment is listed in the figure legends.

### Spinal Cord Cell Dissociation

Mice were anesthetized with 120 mg/kg ketamine and 5mg/kg xylazine and perfused with 15 mL ice-cold Medium A (50 mL 1x HBSS without Ca2+ or Mg2+ (Gibco, 14185052), 750 μL 1M HEPES (Gibco, 15630080), 556 μL 45% glucose (Corning, 25-037-CI)). Lumbar spinal cords were isolated and placed in 2 mL Medium A + 80 μL Dnase I (12,400 units/mL, Sigma, 11284932001) until all samples were dissected. Samples were dounce-homogenized, passed through a 100 μm strainer, washed with 5 mL Medium A, and spun down by centrifugation at 340 x *g* for 7 minutes at 4°C. Supernatant was removed, and pellets were resuspended in 6 mL 25 % Standard Isotonic Percoll (Percoll: GE Healthcare, 17-5445-02 with 10% 10x PBS) in Medium A. The suspension underwent centrifugation for 20 minutes at 950 x *g* at 4°C to remove myelin. Supernatant was discarded then washed with 5 mL Medium A and spun down by centrifugation at 340 x *g* for 7 minutes at 4°C. Cells were resuspended in FACS buffer (5 mM EDTA in 1% BSA in 1x PBS).

### Flow Cytometry

The following antibodies were used for flow cytometry: e450-conjugated anti-CD3 (eBioscience, #48-0031-82), e450-conjugated anti-CD19 (eBioscience, #48-0193-82), PE-Cy7-conjugated anti-CD45 (Biolegend, #103114), APC-conjugated anti-CD11b (Biolegend, #101212). Dead cells were stained with 1:1000 SYTOX Blue (Invitrogen, #S34857). Cell suspension was then spun down at 300 x *g* for 5 minutes at 4°C. The FACS buffer supernatant was suctioned off, and cells were resuspended in fresh FACS buffer. Before staining, all samples were pre-blocked with 1:100 anti-CD16/CD32 (BD Pharmingen, #553142) for 5 minutes at room temperature. Afterward, all antibodies were added to the samples at 1:100 along with 1:1000 Sytox Blue and placed on ice for 30 minutes. Samples were spun down for 5 minutes at 400 x *g* at 4°C, suctioned and replaced with fresh FACS buffer. Suspension was then spun down for 5 minutes at 400 x *g* at 4°C. Supernatant was suctioned off, then cells were again resuspended in FACS buffer and passed through 35 μm filter into polystyrene tubes. Samples were sorted using a BD Aria II (BD Biosciences). Flow cytometry analysis was done using FlowJo v10.6.2. Single cells were gated for live (SYTOX blue negative), CD19-CD3-CD45^mid^ then CD11b^+^ and Cx3CR1-YFP^+^ to isolate microglia from spinal cord.

### Preparation of samples for sequencing

For male samples, whole RNA was isolated from FACS microglia using the Rneasy® Plus Micro kit (Qiagen, Cat No. 74034) then cDNA was synthesized using the SMART-Seq® v4 Ultra® Low Input RNA kit (Takara Bio USA Inc. Cat No. 634888). Libraries were prepared using the Nextera^TM^ DNA Flex Library Prep (Illumina Cat No. 20015828) and sequencing performed by the Stanford Genomics Service Center on a HiSeq4000 using a paired end 2×150bp unstranded protocol. For the female samples, microglia were isolated by FACS into Trizol LS (Invitrogen, Cat No. 10296028) and sent to Azenta Life Sciences (formerly Genewiz) to isolate total RNA using a routine protocol with glycogen added in the precipitation step followed by sequencing on an Illumina HiSeq using paired end 2×150bp unstranded protocol.

### DESeq2 analysis

Raw sequencing data was analyzed using the nf-core/rnaseq v3.5 RNA-Seq analysis pipeline (https://nf-co.re/rnaseq) {Ewels, 2020 #1757} with nf-core helper tools v2.2 (https://github.com/nf-core/tools) {Ewels, 2020 #1757} which was run using STAR and RSEM with isoform (transcript) counts and extensive quality control. The Nextflow v22.01.0 workflow tool (https://github.com/nextflow-io/nextflow) {Di Tommaso, 2017 #1758} facilitated the running of various pipeline tasks across a compute cluster in a very portable manner. FastQC v0.11.9 (https://www.bioinformatics.babraham.ac.uk/projects/fastqc/) {Andrews, 2010 #1761} was used to run quality control checks on raw FASTQ sequence data. The wrapper tool Trim Galore v0.6.7 (https://www.bioinformatics.babraham.ac.uk/projects/trim_galore/) {Krueger, 2021 #1763} that employs Cutadapt v3.4 and FastQC v0.11.9 was run to consistently apply quality and adapter trimming to FASTQ files. Alignment to the GENCODE GRCm38 release M25 reference was done using STAR v2.7.6a and the aligned transcripts were quantitated by RSEM v1.3.1 (http://deweylab.github.io/RSEM/).{Li, 2011 #1759} MultiQC v1.11 (https://multiqc.info/) {Ewels, 2016 #1760} helped to aggregate results from the various bioinformatics analyses across the many samples into a single report. Differentially expressed transcripts were detected from raw transcript counts produced by the STAR-RSEM pipeline using DESeq2 v1.34.0 (https://bioconductor.org/packages/release/bioc/html/DESeq2.html) {Love, 2014 #1762} running in R v4.1.3.

### Weighted gene co-expression network analysis

Weighted gene co-expression network analysis (WGCNA) was used to identify clusters (modules) of highly correlated genes. Uninjured-Resident, Injured-Resident, Uninjured-Repopulated, and Injured-Repopulated replicates from female and male mice were analyzed using WGCNA v1.70.3 package in R v4.1.3.{Langfelder, 2008 #1653} Variance stabilization transformed RNA-Seq expression data was loaded and genes and samples with too many missing values, if any, were removed. Samples were then clustered based on their Euclidean distance to see whether there were any obvious outliers. Traits of interest including Uninjured-Repopulated microglia vs. Uninjured-Resident microglia, Injured-Repopulated microglia vs. Uninjured-Repopulated microglia, Injured-Repopulated microglia vs. Injured-Resident microglia, and Injured-Resident microglia vs. Uninjured-Resident microglia were visualized to see how they related to the sample dendrogram. A step-by-step network and module detection approach was then employed. The soft thresholding power to which co-expression similarity was raised to calculate adjacency was chosen based on the criterion of approximate scale-free topology. Adjacencies were calculated based on the chosen soft thresholding power. To minimize effects of noise and spurious associations, the adjacency was transformed into Topological Overlap Matrix (TOM) following which the corresponding dissimilarity was calculated. Hierarchical clustering was employed to produce a dendrogram of genes. The Dynamic Tree Cut R library that implements novel dynamic branch cutting methods was used to detect clusters (modules) of highly co-expressed genes in the dendrogram. Modules whose expression profiles were very similar were merged after calculating eigengenes of modules and clustering them on their correlation. A cut height of 0.25 corresponding to a correlation of 0.75 was used to merge modules and assign module colors. In the next step modules that were significantly associated with above-mentioned traits of interest such as Uninjured-Repopulated microglia vs. Uninjured-Resident microglia, Injured-Repopulated microglia vs. Uninjured-Repopulated microglia, Injured-Repopulated microglia vs. Injured-Resident microglia, and Injured-Resident microglia vs. Uninjured-Resident microglia were identified by quantifying the module-trait relationships by correlation values and p-values. Next, associations of individual genes with each trait of interest were quantified by defining gene significance (GS) as the absolute value of the correlation between the gene and the trait. For each module, a quantitative measure of module membership (MM) was defined as the correlation of the module eigengene and the gene expression profile which allowed quantification of the similarity of all genes to every module. The GS and MM measures were used to identify genes that have a high significance for each trait of interest as well as high module membership in interesting modules. Finally, a table was created to show gene annotation, module color, gene significance for traits of interest with p-values, and module membership with p-values in all modules.

### Gene ontology and signaling pathway analysis of Gene sets

Gene sets of interest were analyzed via the g:profiler website{Raudvere, 2019 #1815} to identify enriched Gene Ontology Terms and KEGG Pathways based on a threshold of Benjamini-Hochberg FDR > 0.05. Gene Ontology terms were visualized with GO-Figure! Version 1.0.1{Reijnders, 2021 #1816} utilizing the 2021-02-01 release of the Gene Ontology (10.5281/zenodo.2529950) and the 2022-03-02 release of the UniProt GO Associations.{Huntley, 2015 #1817} The pathway analysis was executed through the use of QIAGEN IPA (QIAGEN Inc., https://digitalinsights.qiagen.com/IPA). {Kramer, 2014 #1818}

### Isolation of postmortem human microglia nuclei

Human lumbar spinal cord was removed immediately after organs were removed for transplant from either male or female donor (see Supplemental Table S2). The lumbar segment was further dissected into three smaller segments, and immediately frozen on dry ice and stored at −80°C in Eppendorf or conical tubes. Lumbar spinal cord was then embedded in OCT and sectioned at 80 µm thickness (40-50 total sections per sample was used). Dorsal horn was collected by microdissection and immediately homogenized in Nuclei Lysis buffer on ice (10 mM Tris-HCl pH 8.0, 250 mM sucrose, 25 mM KCl, 5 mM MgCl2, 0.1% Triton-x 100, 0.5% Rnasin® Ribonuclease Inhibitors (Fisher/Promega, PRN2615), 1X Protease inhibitor, 0.1 mM DTT). Samples were then filtered through a 40 µm nylon cell strainer (BD#352340) pre-wet with Nuclei Lysis buffer in a 15 mL conical tube and another 2 mL of lysis buffer was used to rinse the filter. The homogenate was then centrifuged at 900 × *g* for 10 min at 4°C. The resulting pellet was resuspended in 6 mL of sterile 25% Standard Isotonic Percoll (SIP) to remove myelin and centrifuged at 2000 x *g* for 20 min at 4°C without brake. The myelin layer was discarded and the nuclei pellet was resuspended in 1 mL of Staining buffer (1XPBS, 0.8%BSA, 0.5% Rnasin plus). The nuclei suspension was transferred to an Eppendorf tube and centrifuged at 900 × *g* for 10 min at 4°C. Supernatant was discarded and the pellet was resuspended in Staining buffer containing the Human BD Fc Block™ antibody (BD, 564220) and incubated with agitation for 15 min at 4°C in a cold room. Samples were then incubated with rabbit anti-NeuN Alexa Fluor® 647 (1:1000; Abcam, ab190565), rabbit anti-Sox10 Alexa Fluor® 647 (1:5000; Abcam, ab270151), rabbit anti-Sox9 Alexa Fluor® 555 (1:1000; Millipore, AB5535-AF555), and DAPI (1:1000, ThermoFisher, 62248). Samples were incubated with agitation for 30 mins at 4 °C (protected from light) and centrifuged at 900 × *g* for 10 minutes at 4°C. Samples were then resuspended in 1mL Staining buffer and centrifuged at 900 × *g* for 10 minutes at 4°C. Nuclei were resuspended in 400-500 µl of Staining buffer and filtered through the lid of 35μm BD FACS tube before sorting on a BD Aria II (BD Biosciences) machine. DAPI only positive nuclei were sorted away from nuclei co-positive with any other signal to negatively select for and enrich nuclei sample for microglia. Nuclei were then centrifuged at 200 x *g* for 1 min using gentle brake and acceleration and repeated two additional times. Supernatant was removed to leave ∼ 60 µl to resuspend nuclei.

### Single-nuclei RNA sequencing

Nuclei were counted on a hemocytometer and estimated nuclei/µl were calculated for loading onto a 10X Genomic Chromium single cell chip (10x Genomics). Reverse transcription and subsequent library preparations were performed using the Chromium Next GEM Single Cell 3LJ Kit v3 by the Stanford Functional Genomics Facility per manufacturer’s instructions. Samples were then sequenced to an average depth of 97,000-120,000 reads per nuclei on an Illumina NovaSeq6000 sequencer.

### snRNA-seq data processing

Cell Ranger (version 6.0.0) was used for barcode processing, unique molecular identifiers filtering, gene counting, and sample mapping to the reference transcriptome (GRCh38). The filtered UMI count matrix was provided from Cell Ranger as an input in the Seurat workflow (version 4.0).{Stuart, 2019 #1756} Nuclei with less than 200 and greater than 6,000 (for the male sample) or 8,000 (for the female sample) detected genes based on read depth were filtered out to eliminate low quality nuclei and multiplets. Nuclei with more than 5% mitochondrial genes detected were further removed. After filtering, a total of 6,591 nuclei were obtained, 2,782 nuclei for the male sample and 3,809 nuclei for the female sample.

We applied the anchor-based strategy in the Seurat workflow to integrate samples from different batches. To merge two samples, both male and female count matrices were log-normalized in Seurat’s function ‘NormalizeData’ and the top 2,000 shared highly variable genes were found using the function ‘FinderVariableFeatures’ with “selection.method = ‘vst’”. The integration anchors were then identified using the canonical correlation analysis from the function ‘FinderIntegrationAnchors’. Based on the anchors, two datasets were integrated using the function ‘IntegrateData’.

Principal component analysis (PCA) was performed on the integrated dataset. Based on the first 30 principal components (PC), the uniform manifold approximation and projection (UMAP) was used for dimension reduction. A shared nearest neighbor (SNN) graph was built with the first 30 PCs using Seurat’s function ‘FindNeighbors’. The nuclei were clustered using a Louvain algorithm applied by the function ‘FindClusters’.

### Identification of microglia nuclei

Cell types were annotated based on the reference dataset{Habib, 2017 #1752} using Seurat’s label-transferring approach. The identified nuclei clusters were comparable to the unsupervised clustering results under the resolution parameter 0.04. We compared the clusters from reference-based annotation and unsupervised clustering analysis to determine the subset of microglia-specific nuclei.

We found a sub-cluster of the reference-annotated microglia (cluster 4 from the unsupervised clustering results) highly expressed T cell signature genes (*THEMIS*, *SKAP1*) (data not shown). These nuclei were excluded in further analysis. The final microglia nuclei were selected as the intersection of the reference-annotated microglia cells and the nuclei from clusters 1 and 3 in unsupervised clustering analysis (boxed in Figure 6c). It was confirmed that microglia marker genes (*ITGAM*, *CTSS*, *CSF1R*, *C1QA*) were highly expressed in these clusters (boxed in Figure 6c and 6d). Using these criteria, 2,316 nuclei were classified as microglia and isolated for further analysis.

### Sub-cluster annotation for microglia nuclei

Using the isolated microglia nuclei, we applied the unsupervised clustering analysis under a series of resolution parameters to investigate biological meaningful subclusters. Microglia were clustered using resolution 0.06 to form 4 clusters. Based on differential gene expression patterns, cluster 0 was further sub-clustered into three additional clusters (cluster 2, 3 and 6). After this regrouping, we ended with 6 total microglia nuclei subclusters (Figure 6e). Differential expression analysis was performed from the function ‘FindAllMarkers’ in the Seurat package with filtering parameters ‘min.pct = 0.25’ and ‘logfc.threshold = 0.25’.

### Study Approval

All procedures were approved by the Stanford University Administrative Panel on Laboratory Animal Care and the Veterans Affairs Palo Alto Health Care System Institutional Animal Care and Use Committee in accordance with American Veterinary Medical Association guidelines and the International Association for the Study of Pain. Human post-mortem spinal cord was obtained in collaboration with Donor Network West and received Institutional Review Board exemption.

## Supplemental Information Titles and Legends

**Fig. S1. Microglial depletion alone has no behavioral impact but depletion in the acute phase results in partial improvement in pain sensitivity and peripheral signs of inflammation**

Microglial-depletion alone does not affect baseline a) weight bearing, b) paw thickness, or c) paw temperature in male or female Cx3CR1-creERT2-eYFP;R26-iDTRLSL mice (n = 5-6/sex). Data are represented as mean ± SEM. d) Schematic of experimental timeline for acute phase (Day 0) depletion of microglia. e) Mechanical threshold is decreased at the time of cast removal (3 weeks) in all groups, however, microglial depletion results in progressive, partial improvement in mechanical sensitivity in both males and females out to 9 weeks post-injury. n = 5-15 per group per sex. **p< 0.01, ***p<0.001, ****p<0.0001 by two-way ANOVA with Bonferroni’s post-hoc test. Thermal latency on a 52.5°C hot plate is not significantly improved in males (f) but is in females (g) after microglial depletion. Injured Microglia-depleted mice improved the amount of weight placed on the injured paw (h, males; i, females), exhibited less edema (j, males; k, females) and less temperature change (l, males; m, females) compared to injured mice without microglia depletion. n = 4-8/sex/group. *p<0.05, **p< 0.01, ***p<0.001, ****p<0.0001 by unpaired t-test. Data are represented as mean ± SEM.

**Fig. S2. Fluorescence-Activated Cell Sorting (FACS) of spinal cord microglia and principal component analysis of transcriptome data from each sample**

a) Gating strategy for sorting microglia from spinal cord. Live (SYTOX blue negative), CD19-CD3-CD45mid then CD11b+ and Cx3CR1-YFP+. SSC-A: side scatter area; FSC-A: forward scatter area; SSC-W: side scatter width; SSC-H: side scatter height; FSC-W: forward scatter width. b) To confirm the specificity of our isolation method for CNS microglia, we used the publicly available ImmGen “My GeneSet” program to cross-reference genes in our male RNAseq dataset with the ImmGen database of over 50 cell populations. The top 100 genes expressed by log2(cpm+1) in our datasets clearly defined a microglia-specific signature that could be differentiated from similar myeloid-lineage cells including spleen macrophages and Ly6C_hi and Ly6C_lo blood monocytes. c) Principal component analysis (PCA) of each female sample’s transcriptomic data. Analysis maps each sample in two-dimensional space based on the two principal components responsible for the most variance in the samples. Female PC1, accounting for 48% of the variance, splits the samples based on injured mice or uninjured mice. d) PCA of each male sample’s transcriptomic data. Male PC1, accounting for 45% of the variance, shows distinct separation between the groups and especially splits resident microglia and repopulated microglia.

**Fig. S3. Volcano plots of DEseq2 data for the other three comparisons**

a) Volcano plots for each sex of the comparison of injured mice with resident microglia vs uninjured mice with resident microglia. Each transcript from the deseq2 analysis is plotted as a point on the graph by log2(fold change) and −log10(adjusted p-value). Cutoffs set at x=+/-1 and y=1.3 for both sexes. b) Same style of volcano plots for the comparison of uninjured mice with repopulated microglia versus uninjured mice with resident microglia. Cutoffs set at x=+/-1.5 for females, x=+/-1 for males and y=1.3 for both sexes. c) Same style of volcano plots for the comparison of injured mice with repopulated microglia versus uninjured mice with repopulated microglia. Cutoffs set at x=+/-1.5 for females, x=+/-1 for males and y=1.3 for both sexes.

**Fig. S4. WGCNA cluster dendrograms for all transcripts**

Hierarchical clustering dendrograms visualize the transcripts analyzed via WGCNA, distributed based on transcript-to-transcript adjacency. Transcripts are grouped into modules based on a dynamic tree cut algorithm that identifies branches of the dendrogram to cluster together. Each original module is represented by a color in the top half of the color row below the dendrogram. Lastly, modules with high similarity are merged into dynamic modules depicted in the bottom half of the color row.

**Fig. S5. Transcriptome shift in males and females of the Injured-Repopulated vs Injured-Resident comparison**

a) The transcripts that are in both a male and female module of interest are mapped in the same way. Pearson’s correlation value is R^2^=0.6802, showing that the modules of interest are capturing a transcriptome shift that is shared in males and females. b) All transcripts that were in both the male and female DESeq2 data were plotted by x=log2(fold change) in males and y=log2(fold change) in females. Proximity to the x=y line on the plot indicates similar expression changes in males and females. Pearson’s correlation value of R^2^=0.0714.

**Fig. S6. Key genes expressed in a human microglia cluster-specific manner**

a) MKI67, known as Ki-67, and NDC80 is expressed solely in cluster 5, denoted as a proliferative microglia cluster. b) TLR2, CYBB (NOX2), and AC083837.1 highly expressed in cluster 4, denoted as an ‘reactive’ microglia cluster. c) ITGAX, IGF1R, TGFBI, and SAMD9L expressed significantly in cluster 1, denoted as a homeostatic microglia cluster. p-values from differential cluster-expression analysis listed in parentheses under gene name for each respective cluster in panels.

**Table S1.** Differentially expressed transcripts (DETs) and differentially expressed genes (DEGs) for all comparisons

**Table S2.** Characteristics of human donors

**Table S3.** Number of microglial nuclei per sub-cluster

**Table S4.** Overlapping mouse genes of interest from transcriptomic analysis via DESeq2 and WGCNA Integration of the two primary analyses applied to the mouse transcriptome data, DESeq2 and WGCNA, generated a list of genes of interest for further study. Genes from our comparison of interest, Injured-Repopulated microglia vs Injured-Resident microglia, were broken down into three categories: genes significantly expressed in both sexes, genes specific to females, and genes specific to males.

Extended Data 1. Differentially expressed transcripts and genes for each comparison in both sexes

Extended Data 2. Gene Ontology terms enriched by differentially expressed genes for each comparison in females

Extended Data 3. Gene Ontology terms enriched by differentially expressed genes for each comparison in males

Extended Data 4. Gene Ontology terms enriched by differentially expressed genes that overlapped between sexes for each comparison

Extended Data 5. Correlation coefficients and p-values for the module-trait heatmaps Extended Data 6. Distribution of differentially expressed transcripts in WGCNA modules

Extended Data 7. Ingenuity pathway analysis of full list of genes in modules of interest and the top 20 genes by gene significance and module membership in each module

Extended Data 8. Differentially expressed genes in microglia subclusters of human single nuclei RNA-seq dataset

Extended Data 9. Gene Ontology terms enriched by differentially expressed genes microglia subclusters of human single nuclei RNA-seq dataset

