## Supplemental tables and figures for "Newly repopulated spinal cord microglia exhibit a unique transcriptome and correlate with pain resolution"

### Supplemental Figure 1

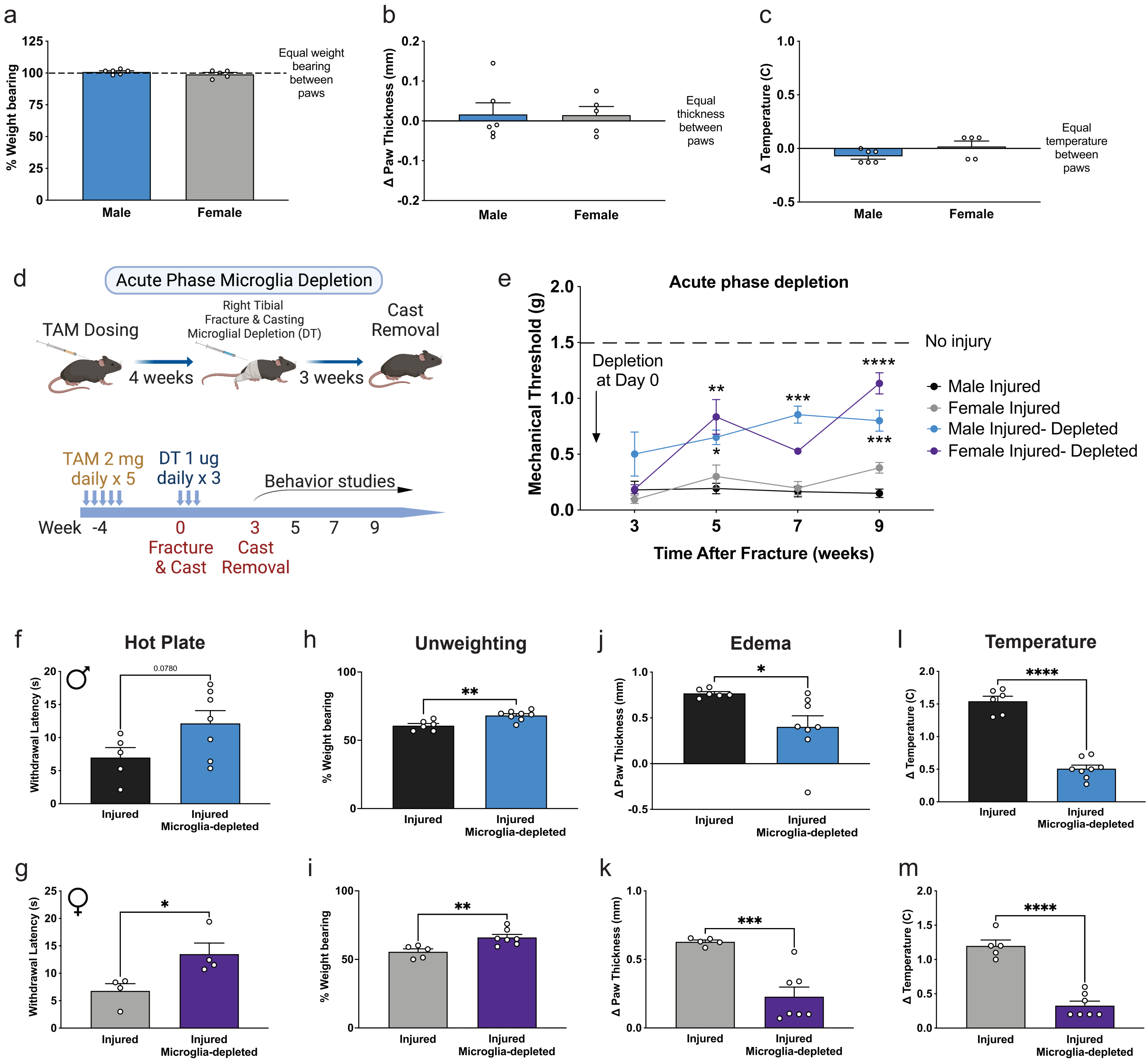

**Fig. S1. Microglial depletion alone has no behavioral impact but depletion in the acute phase results in partial improvement in pain sensitivity and peripheral signs of inflammation**

Microglial-depletion alone does not affect baseline a) weight bearing, b) paw thickness, or c) paw temperature in male or female Cx3CR1-creERT2-eYFP;R26-iDTRLSL mice (n = 5-6/sex). Data are represented as mean ± SEM. d) Schematic of experimental timeline for acute phase (Day 0) depletion of microglia. e) Mechanical threshold is decreased at the time of cast removal (3 weeks) in all groups, however, microglial depletion results in progressive, partial improvement in mechanical sensitivity in both males and females out to 9 weeks post-injury. n = 5-15 per group per sex. **$p < 0.01$, ***$p<0.001$, ****$p<0.0001$ by two-way ANOVA with Bonferroni's post-hoc test. Thermal latency on a 52.5ºC hot plate is not significantly improved in males (f) but is in females (g) after microglial depletion. Injured Microglia-depleted mice improved the amount of weight placed on the injured paw (h, males; i, females), exhibited less edema (j, males; k, females) and less temperature change (l, males; m, females) compared to injured mice without microglia depletion. n = 4-8/sex/group. *$p<0.05$, **$p< 0.01$, ***$p<0.001$, ****$p<0.0001$ by unpaired t-test. Data are represented as mean ± SEM.

Supplemental Figure 2

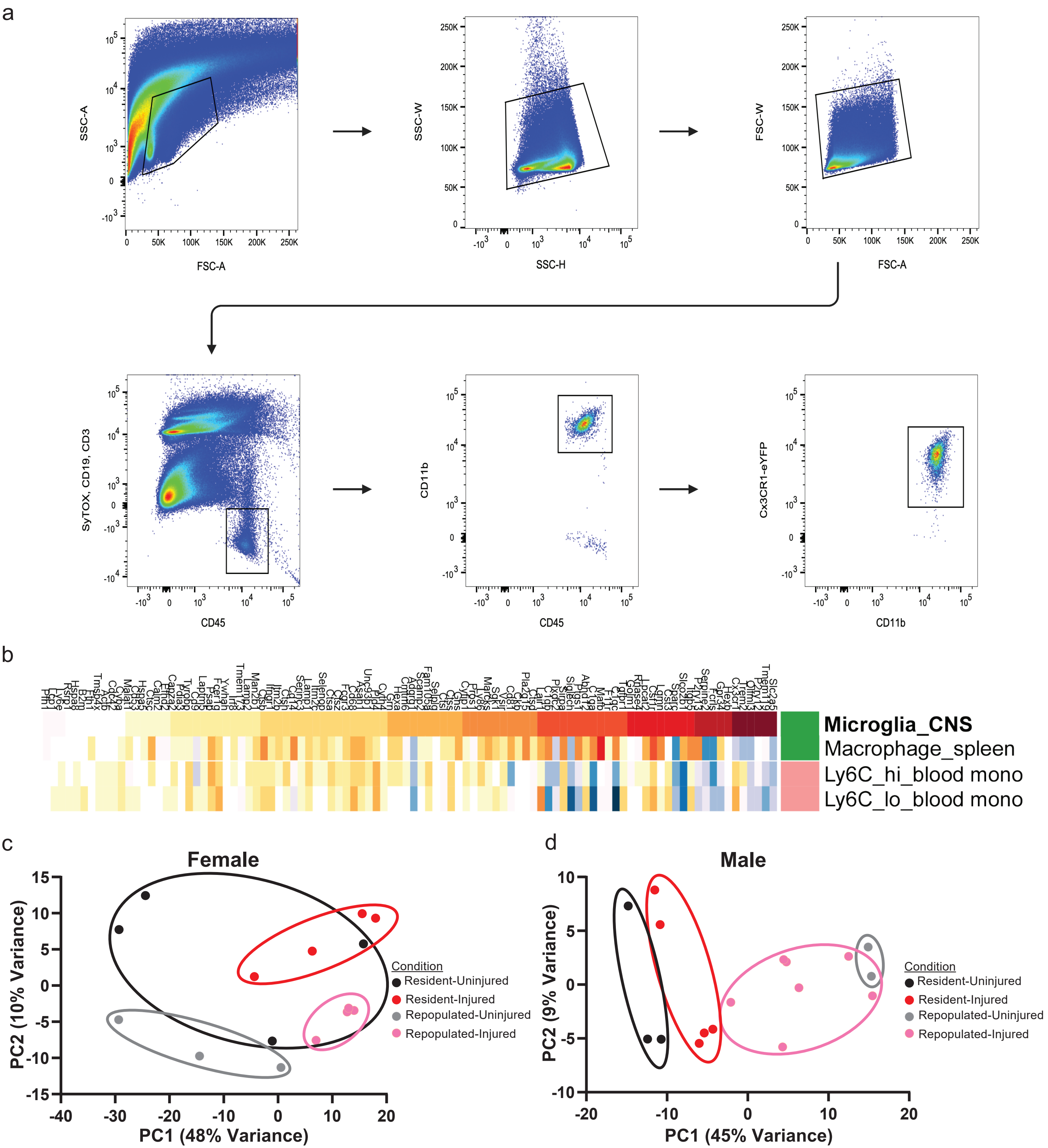

**Fig. S2. Fluorescence-Activated Cell Sorting (FACS) of spinal cord microglia and principal component analysis of transcriptome data from each sample**

a) Gating strategy for sorting microglia from spinal cord. Live (SYTOX blue negative), CD19- CD3- CD45mid then CD11b+ and Cx3CR1-YFP+. SSC-A: side scatter area; FSC-A: forward scatter area; SSC-W: side scatter width; SSC-H: side scatter height; FSC-W: forward scatter width. b) To confirm the specificity of our isolation method for CNS microglia, we used the publicly available ImmGen "My GeneSet" program to cross-reference genes in our male RNAseq dataset with the ImmGen database of over 50 cell populations. The top 100 genes expressed by log2(cpm+1) in our datasets clearly defined a microglia-specific signature that could be differentiated from similar myeloid-lineage cells including spleen macrophages and Ly6C_hi and Ly6C_lo blood monocytes. c) Principal component analysis (PCA) of each female sample's transcriptomic data. Analysis maps each sample in two-dimensional space based on the two principal components responsible for the most variance in the samples. Female PC1, accounting for 48% of the variance, splits the samples based on injured mice or uninjured mice. d) PCA of each male sample's transcriptomic data. Male PC1, accounting for 45% of the variance, shows distinct separation between the groups and especially splits resident microglia and repopulated microglia.

Supplemental Figure 3

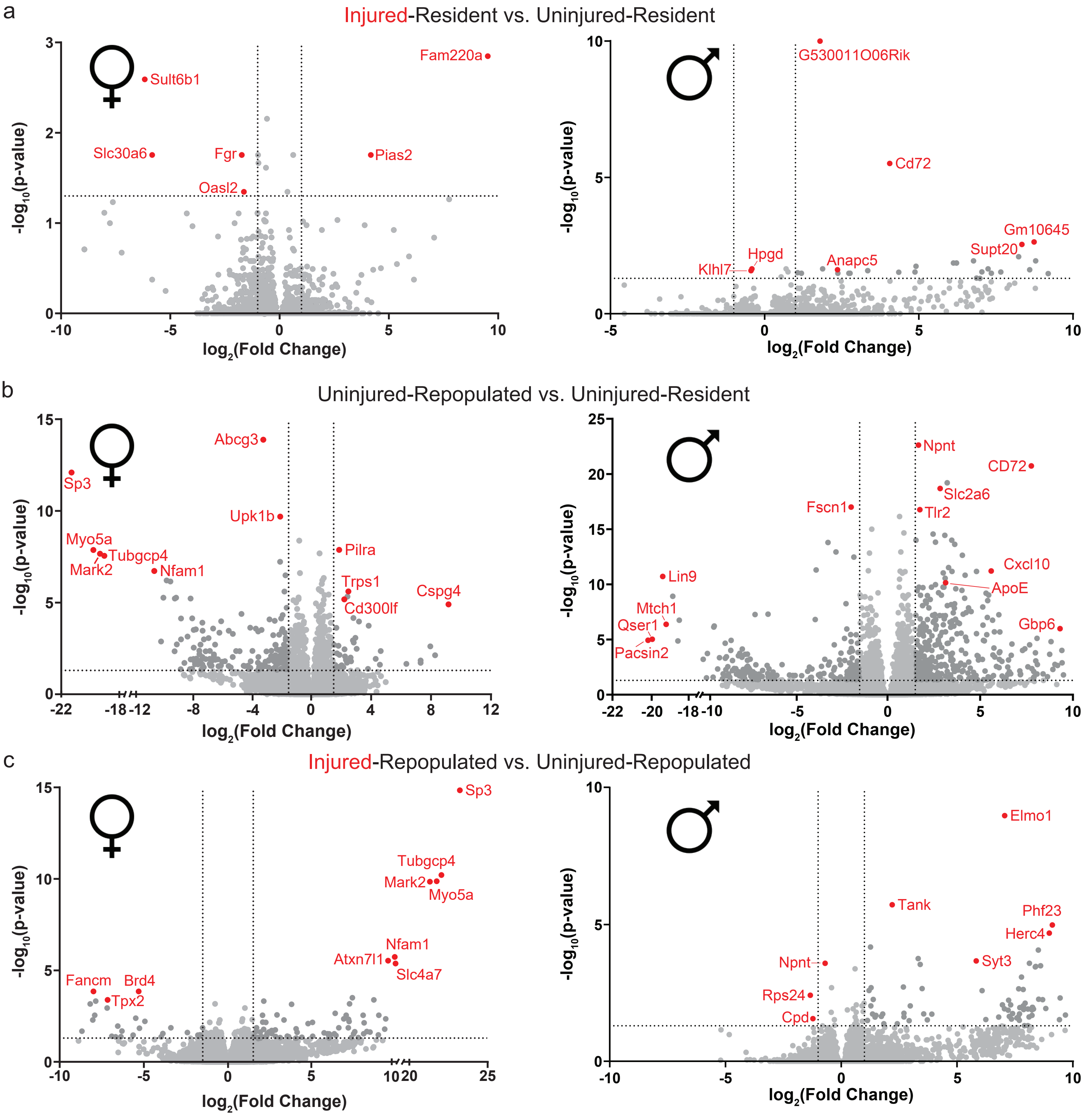

**Fig. S3. Volcano plots of DEseq2 data for the other three comparisons**

a) Volcano plots for each sex of the comparison of injured mice with resident microglia vs uninjured mice with resident microglia. Each transcript from the deseq2 analysis is plotted as a point on the graph by log2(fold change) and -log10(adjusted p-value). Cutoffs set at $|x| > 1$ and $y > 1.3$ for both sexes. b) Same style of volcano plots for the comparison of uninjured mice with repopulated microglia versus uninjured mice with resident microglia. Thresholds are set at $|x| > 1.5$ for females, $|x| > 1$ for males and $y > 1.3$ for both sexes. c) Same style of volcano plots for the comparison of injured mice with repopulated microglia versus uninjured mice with repopulated microglia. Cutoffs set at $|x| > 1.5$ for females, $|x| > 1$ for males and $y > 1.3$ for both sexes.

Supplemental Figure 4

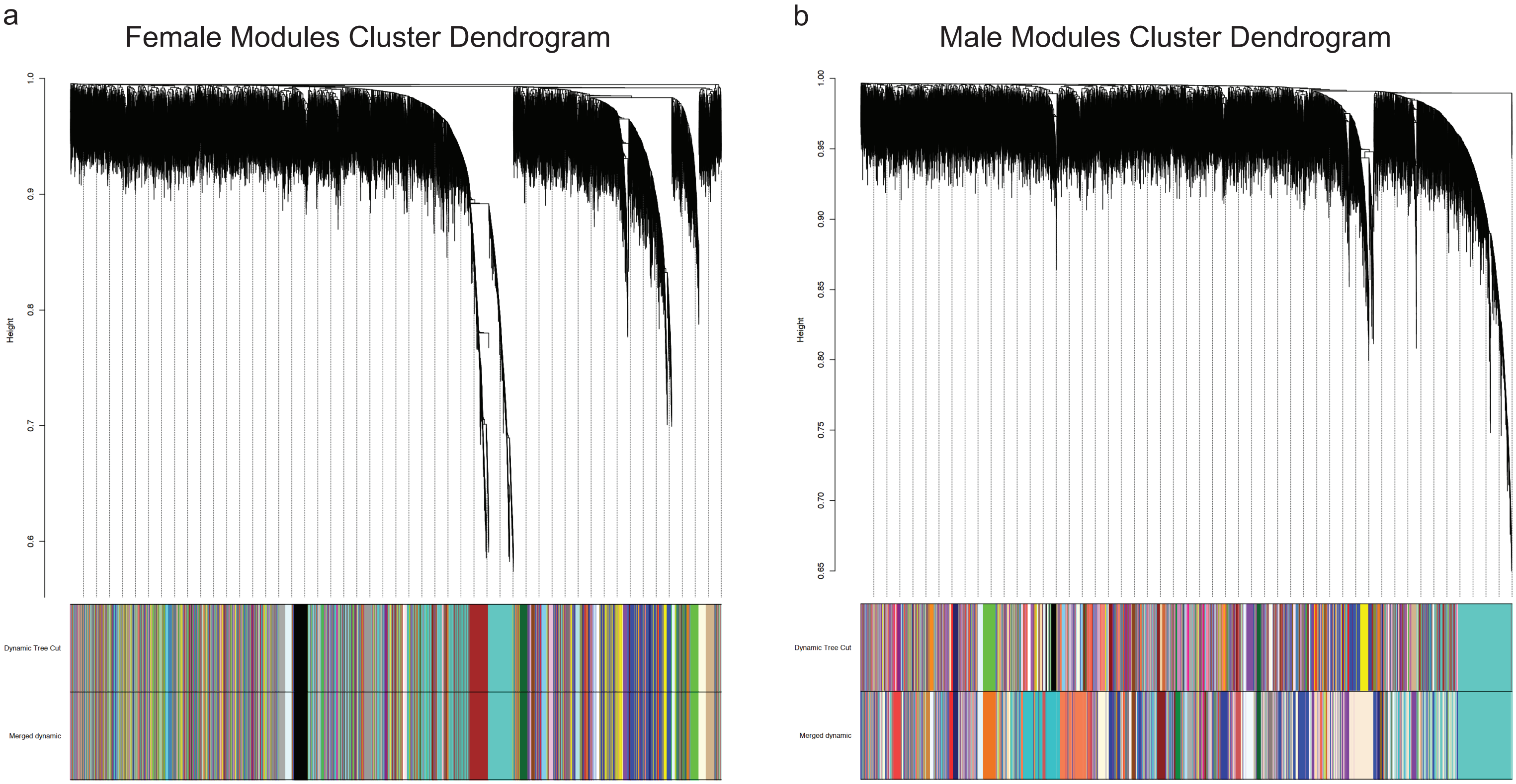

**Fig. S4. WGCNA cluster dendrograms for all transcripts**

Hierarchical clustering dendrograms visualize the transcripts analyzed via WGCNA, distributed based on transcript-to-transcript adjacency. Transcripts are grouped into modules based on a dynamic tree cut algorithm that identifies branches of the dendrogram to cluster together. Each original module is represented by a color in the top half of the color row below the dendrogram. Lastly, modules with high similarity are merged into dynamic modules depicted in the bottom half of the color row.

### Supplemental Figure 5

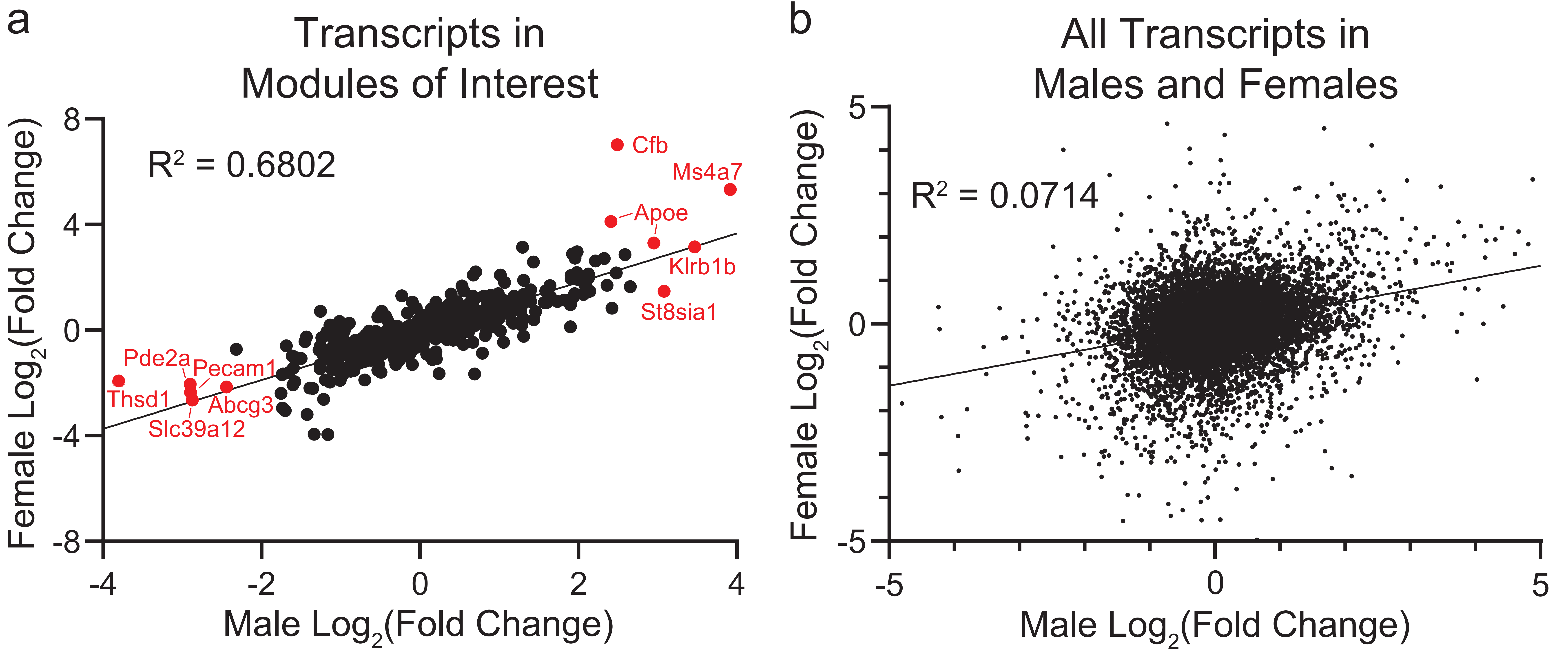

**Fig. S5. Transcriptome shift in males and females of the Injured-Repopulated vs Injured-Resident comparison**

a) The transcripts that are in both a male and female module of interest are mapped in the same way. Pearson's correlation value is $R^2$=0.6802, showing that the modules of interest are capturing a transcriptome shift that is shared in males and females. b) All transcripts that were in both the male and female DESeq2 data were plotted by x=log2(fold change) in males and y=log2(fold change) in females. Proximity to the x=y line on the plot indicates similar expression changes in males and females. Pearson's correlation value of $R^2$=0.0714.

Supplemental Figure 6

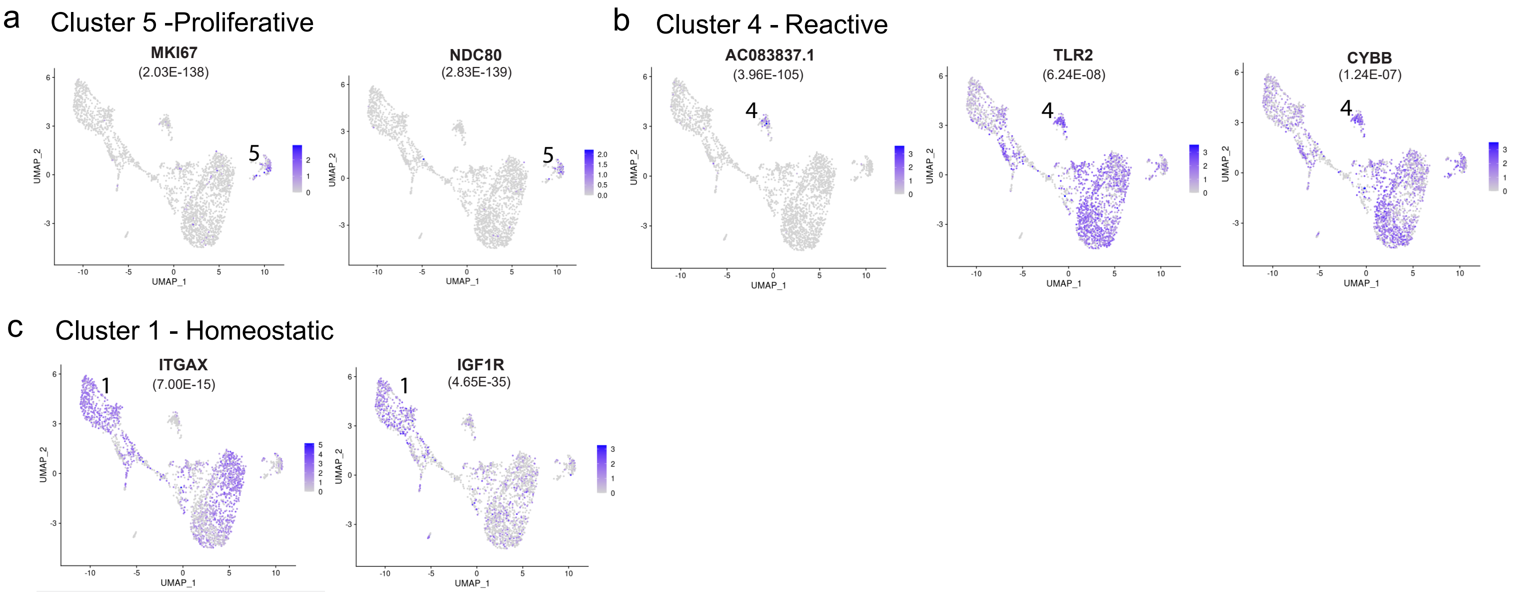

**Fig. S6. Key genes expressed in a human microglia cluster-specific manner**

a) MKI67, known as Ki-67, and NDC80 is expressed solely in cluster 5, denoted as a proliferative microglia cluster. b) TLR2, CYBB (NOX2), and AC083837.1 highly expressed in cluster 4, denoted as an 'reactive' microglia cluster. c) ITGAX and IGF1R expressed significantly in cluster 1, denoted as a homeostatic microglia cluster. p-values from differential cluster-expression analysis listed in parentheses under gene name for each respective cluster in panels.

#### Supplementary Table 1: DETs and DEGs for all comparisons

| | Injured-Resident vs Uninjured-Resident | | | |
|---|---|---|---|---|
| | Transcripts Up | Genes Up | Transcripts Down | Genes Down |
| Male | 32 | 30 | 2 | 2 |
| Female | 4 | 4 | 9 | 8 |
| Overlap | 0 | 0 | 0 | 0 |

| | Uninjured-Repopulated vs Uninjured-Resident | | | |
|---|---|---|---|---|
| | Transcripts Up | Genes Up | Transcripts Down | Genes Down |
| Male | 710 | 571 | 539 | 494 |
| Female | 306 | 251 | 399 | 375 |
| Overlap | 110 | 99 | 80 | 72 |

| | Injured-Repopulated vs Uninjured-Repopulated | | | |
|---|---|---|---|---|
| | Transcripts Up | Genes Up | Transcripts Down | Genes Down |
| Male | 93 | 92 | 7 | 7 |
| Female | 108 | 104 | 61 | 60 |
| Overlap | 1 | 1 | 0 | 0 |

| | Injured-Repopulated vs Injured-Resident | | | |
|---|---|---|---|---|
| | Transcripts Up | Genes Up | Transcripts Down | Genes Down |
| Male | 503 | 412 | 305 | 283 |
| Female | 354 | 293 | 361 | 338 |
| Overlap | 132 | 112 | 75 | 72 |

**Table S1. Differentially expressed transcripts (DETs) and differentially expressed genes (DEGs) for all comparisons**

**Supplementary Table S2: Characteristics of human donors**

| Sex | Age | Cause of Death | History of chronic pain | Ethnicity |
|---|---|---|---|---|
| Male | 43 years old | Anoxia, cardiovascular | None reported | White |
| Female | 49 years old | Anoxia, cardiovascular | None reported | Asian |

**Table S2. Characteristics of human donors**

#### Supplementary Table S3: Number of microglial nuclei per sub-cluster

| Cluster # | Microglia Nuclei count |
|---|---|
| 1 | 730 |
| 2 | 857 |
| 3 | 470 |
| 4 | 81 |
| 5 | 81 |
| 6 | 97 |
| **Total** | **2316** |

**Table S3. Number of microglial nuclei per sub-cluster**

| WGCNA-Filtered Deseq2-based Genes of Interest: Top 30 Genes | | | | | | |
|---|---|---|---|---|---|---|
| Overlap Genes | | | Female-Specific Genes | | Male-Specific Genes | |
| Gene Name | Female FC | Male FC | Gene Name | Female FC | Gene Name | Male FC |
| **Serpinf1** | **-3.0525** | **-1.7013** | Rnaset2b | -9.9720 | Samd9l | 2.4356 |
| **Upk1b** | **-2.0720** | **-1.6003** | Nrp1 | 0.8624 | Ccl2 | 2.0273 |
| **Pilra** | **2.0760** | **1.9069** | Clec12a | 5.7266 | Cxcl13 | 4.1689 |
| Slc2a6 | 1.0829 | 1.7836 | Pde2a | -8.1132 | Khdrbs3 | -2.3230 |
| **Itm2b** | **0.3431** | **0.4049** | Thsd1 | -3.9482 | Fgl2 | 1.7580 |
| **Apoe** | **2.6161** | **2.2157** | Lamb2 | -1.5285 | Ccl12 | 1.5745 |
| Tlr2 | 0.7960 | 1.0348 | Tom1l1 | -3.9388 | Ifit1 | 2.1235 |
| **Scimp** | **1.0901** | **1.7374** | Apobec1 | 1.6100 | Usp18 | 1.9467 |
| **Tmem204** | **-1.6898** | **-1.7388** | Rassf4 | 0.5443 | Naglu | 0.4174 |
| **Ctss** | **0.5801** | **0.5524** | Carmil1 | 3.5239 | Irf7 | 2.9848 |
| **Tnfsf8** | **1.5264** | **1.3974** | Cd300lf | 1.9659 | Ifi213 | 3.6391 |
| Cd72 | 1.6603 | 2.9241 | Trps1 | 2.0721 | Cxcl10 | 2.8610 |
| Oas3 | 2.2575 | 4.0871 | | | Ifi44 | 3.5638 |
| Fscn1 | -0.9490 | -1.0703 | | | Ccl5 | 3.2766 |
| **Gimap6** | **-1.8851** | **-1.1918** | | | Gm2a | 0.4509 |
| **Slc11a1** | **0.4766** | **0.4743** | | | Lamp1 | 0.3881 |
| **Lag3** | **1.0976** | **0.9253** | | | Treml2 | 2.2377 |
| **Abcg3** | **-2.1486** | **-2.4404** | | | Rsad2 | 2.4560 |
| **Ecscr** | **-0.9088** | **-0.7937** | | | Oas1a | 1.6807 |
| H2-K1 | 0.9833 | 1.4478 | | | Npnt | 0.6566 |
| **Ctsh** | **0.4367** | **0.4162** | | | Slfn5 | 2.1340 |
| Scpep1 | 0.3662 | 0.6112 | | | Ifit3 | 2.1351 |
| Axl | 1.4376 | 1.8007 | | | Phf11a | 1.9342 |
| Ifi211 | 1.4120 | 3.0868 | | | Ifi204 | 1.6425 |
| **Ms4a7** | **5.3212** | **3.9165** | | | Atp10a | 3.0844 |
| **Rtn1** | **-0.6677** | **-0.6115** | | | Oas2 | 4.3090 |
| **Hcar2** | **0.9187** | **1.2809** | | | Rnf213 | 1.4956 |
| H2-Q7 | 1.7796 | 2.1807 | | | St8sia1 | 3.0809 |
| **Sox4** | **-1.1414** | **-0.8378** | | | Cst7 | 1.5009 |
| **Man2b1** | **0.3648** | **0.4606** | | | | |

**Table S4. Overlapping mouse genes of interest from transcriptomic analysis via DESeq2 and WGCNA**

Integration of the two primary analyses applied to the mouse transcriptome data, DESeq2 and WGCNA, generated a list of genes of interest for further study. Genes from our comparison of interest, Injured-Repopulated microglia vs Injured-Resident microglia, were broken down into three categories: genes significantly expressed in both sexes, genes specific to females, and genes specific to males.

**Extended Data 1. Differentially expressed transcripts and genes for each comparison in both sexes**

**Extended Data 2. Gene Ontology terms enriched by differentially expressed genes for each comparison in females**

**Extended Data 3. Gene Ontology terms enriched by differentially expressed genes for each comparison in males**

**Extended Data 4. Gene Ontology terms enriched by differentially expressed genes that overlapped between sexes for each comparison**

**Extended Data 5. Correlation coefficients and p-values for the module-trait heatmaps**

**Extended Data 6. Distribution of differentially expressed transcripts in WGCNA modules**

**Extended Data 7. Ingenuity pathway analysis of full list of genes in modules of interest and the top 20 genes by gene significance and module membership in each module**

**Extended Data 8. Differentially expressed genes in microglia subclusters of human single nuclei RNA-seq dataset**

**Extended Data 9. Gene Ontology terms enriched by differentially expressed genes microglia subclusters of human single nuclei RNA-seq dataset**
